# *let-7* controls the transition to adulthood by releasing select transcriptional regulators from repression by LIN41

**DOI:** 10.1101/460352

**Authors:** Florian Aeschimann, Anca Neagu, Magdalene Rausch, Helge Großhans

## Abstract

Development of multicellular organisms relies on faithful temporal control of cell fates. In *Caenorhabditis elegans*, the heterochronic pathway governs temporal patterning of somatic cells. This function may be phylogenetically conserved as several heterochronic genes have mammalian orthologues, and as the heterochronic *let-7* miRNA and its regulator LIN28 appear to time the onset of puberty in mice and humans. Here, we have investigated how *let-7* promotes the transition to adulthood in *C. elegans*. We find that *let-7* controls each of three relevant processes, namely male and female sexual organ morphogenesis as well as changes in skin progenitor cell fates, through the same single target, *lin-41*. LIN41 in turn silences two pairs of targets post-transcriptionally, by binding and silencing their mRNAs. The EGR-type transcription factor LIN-29a and its co-factor, the NAB1/2 orthologous MAB-10, mediate control of progenitor cell fates and vulval integrity. By contrast, male tail development depends on regulation of the DM domain-containing transcription factors DMD-3 and MAB-3. Our results provide mechanistic insight into an exemplary temporal patterning pathway, demonstrate that *let-7* – LIN41 function as a versatile regulatory module that can be connected to different outputs, and reveal how several levels of post-transcriptional regulation ultimately achieve effects through controlling transcriptional outputs.

## INTRODUCTION

Development of multicellular organisms requires faithful control of cell fates in both space and time. Although our knowledge of temporal control mechanisms lacks a detailed mechanistic understanding rivalling that of spatial patterning, fundamental and conserved regulators of developmental timing in animals have been identified. Specifically, the miRNA *let-7* and its regulator LIN28 control not only transitions between self-renewal and differentiation programs of stem and progenitor cells in various contexts, but also sexual maturation of organisms as distinct as *C. elegans* and mammals (Corre et al., 2016; Faunes and Larrain, 2016).

The realization that temporal patterning in highly distinct organisms may extend beyond shared general principles to specific molecular mechanisms has emphasized the utility of studying developmental timing in *C. elegans*, where temporal patterning is highly stereotyped. Characterization of *C. elegans* mutations that change timing of specific developmental events relative to other developmental events has led to the discovery of the heterochronic pathway that controls temporal somatic cell fates (Ambros, 1989; Ambros and Horvitz, 1984). Among the heterochronic genes, the *let-7* miRNA stands out through its perfect sequence conservation from worm to human, and the fact that it is the only heterochronic gene that is absolutely essential for worm survival. The heterochronic pathway genes in general, and *let-7* in particular, control the transition from a juvenile to an adult animal. Among other events, this transition involves formation of mature sexual organs, i.e., the hermaphrodite vulva and the male tail, respectively. Aberrant morphogenesis of female genitalia appears to be the reason why hermaphrodites with dysfunctional *let-7* rupture through the vulva and die (Ecsedi et al., 2015). More immediately reflecting a role of *let-7* in temporal patterning, impaired *let-7* function delays the onset of the cell retraction events that shape the male tail into a functional sexual organ (Del Rio-Albrechtsen et al., 2006).

In addition to morphogenetically visible changes, transition of *C. elegans* larvae to adulthood also affects specific cell fates. Thus, whereas epidermal progenitor cells called seam cells divide asymmetrically in each larval stage, they exit the cell cycle upon transition into adulthood (Sulston and Horvitz, 1977). As in certain mammalian stem and progenitor cells (Büssing et al., 2008), *let-7* promotes this change in cell fate. When *let-7* activity is lost, the switch to the adult fate fails so that seam cells keep their larval fate and continue to divide in adult animals (Reinhart et al., 2000).

The molecular basis of the *let-7* mutant phenotypes has only begun to emerge. MicroRNAs silence target mRNAs post-transcriptionally by binding short stretches of complementary sequence, typically located in the 3’ untranslated region (3’ UTR). The low information content of these short sequences means that miRNAs can potentially bind many mRNAs, and many targets of *let-7* have indeed been reported (Abrahante et al., 2003; Andachi, 2008; Großhans et al., 2005; Hunter et al., 2013; Jovanovic et al., 2010; Lin et al., 2003). However, using a genetic strategy that permitted uncoupling of endogenous mRNAs from *let-7*-mediated repression, we could recently demonstrate that the vulval bursting phenotype observed in *let-7* mutant animals relies on only a single target, *lin-41* (Ecsedi et al., 2015). The downregulation of LIN41 through *let-7* during the last larval stages is thus essential for survival. Whether the other *let-7* functions; i.e., control of seam cell self-renewal and male tail development, rely similarly on only one target, and whether this would be *lin-41* or some other transcript, has been unknown.

Here, we demonstrate that the known major functions of *let-7*, control of male and female sexual organ morphogenesis and seam cell fates, all occur through regulation of LIN41 alone. [We refer to the protein by its generic name LIN41 instead of the hyphenated worm-specific LIN-41 to reflect its phylogenetic conservation.] Understanding how sustained accumulation of LIN41 can account for the *let-7* mutant phenotypes requires knowledge of its molecular function, which has been unclear. LIN41 carries features of an RNA-binding protein and an E3 ubiquitin ligase and was also described to have a function as a structural protein (reviewed in (Ecsedi and Großhans, 2013)). The ubiquitin ligase activity was previously shown to mediate some physiological and cellular functions of the mammalian TRIM71/LIN41 protein (Chen et al., 2012; Rybak et al., 2009). However, the RING domain, characteristic of this type of E3 ligases, appears poorly conserved in *C. elegans* LIN41 and dispensable for its heterochronic function (Tocchini et al., 2014).

Recently, we showed that LIN41 can post-transcriptionally regulate mRNAs by direct binding to hairpin structures in their 5’ or 3’ UTRs (Aeschimann et al., 2017; Kumari et al., 2018). We identified and validated four mRNA targets that are silenced by direct binding of LIN41 (Aeschimann et al., 2017). Each of these four LIN41 targets, encoding the transcription factors LIN-29a, DMD-3 and MAB-3, and the transcription cofactor MAB-10, has previously ascribed heterochronic functions (Ambros and Horvitz, 1984; Harris and Horvitz, 2011; Mason et al., 2008). However, most described mutant phenotypes are only partially overlapping with those reported for animals lacking *let-7* or animals with increased LIN41 activity (Del Rio-Albrechtsen et al., 2006; Ecsedi et al., 2015; Euling et al., 1999; Hodgkin, 1983; Mason et al., 2008). Moreover, transcriptome-wide analyses revealed that LIN41 interacted with hundreds of mRNAs in both *C. elegans* and mammalian cells (Kumari et al., 2018; Loedige et al., 2013; Tsukamoto et al., 2017). Thus, whether and to what extent individual LIN41 targets or combinations thereof can explain *let-7* mutant phenotypes awaits clarification.

Here, we have comprehensively addressed how *let-7* and LIN41 control temporal pattering in different tissues. We demonstrate that while *let-7* controls male and female sexual organ morphogenesis and seam cell fates through regulation of LIN41 alone, LIN41 in turn silences mRNAs that encode distinct pairs of effector proteins to execute these functions. Thus, LIN-29a and MAB-10 jointly control seam cell fates and vulval morphogenesis, and we resolve previously observed discrepancies in phenotypes concerning vulval-uterine development by showing that *let-7*-independent accumulation of LIN-29a in the anchor cell is important for uterine development. By contrast, *mab-3* and *dmd-3* are direct targets of LIN41 in male tail morphogenesis, silenced through their 3’UTRs. Indeed, we show that *let-7* – LIN41, through MAB-3 and DMD-3, play a fundamentally essential role for cell retraction events during male tail morphogenesis.

Our results highlight how the *let-7 –* LIN41 module is used as a versatile regulatory module in development. They extend our mechanistic and conceptual understanding of a paradigmatic temporal patterning pathway and reveal how several levels of post-transcriptional regulation promote transition into adulthood through controlling transcriptional programs.

## RESULTS

### LIN41 is the single key target of let-7 for three distinct developmental functions

We previously showed that regulation of LIN41 alone accounted for the function of *let-7* in *C. elegans* vulval development (Ecsedi et al., 2015). Given this unexpected result, we wondered whether other known functions of *let-7* were also explained by a single target. To explore this, we analyzed a set of mutants that enabled testing whether a certain phenotype is caused by failed *let-7*-mediated repression of *lin-41* before the worms reach adulthood (Figure 1A). First, we studied *let-7* mutant phenotypes using *let-7(n2853)* animals. These worms harbor a G-to-A point mutation in the *let-7* seed sequence, preventing *let-7* activity at 25 °C (Reinhart et al., 2000) and thus upregulating all *let-7* targets including *lin-41*. For simplicity, we refer to this allele as *let-7(PM)* for *let-7* point mutation. Second, we tested if preventing *let-7*-mediated silencing of only *lin-41* is sufficient to recapitulate a phenotype. To this end, we used *lin-41(xe8)* animals (Ecsedi et al., 2015) that lack a segment of the *lin-41* 3’UTR carrying the two functional *let-7* target sites. Because these sites were previously referred to as *let-7* complementary sites (LCSs) (Vella et al., 2004), we refer to *lin-41(xe8)* in the following as *lin-41(∆LCS)*. Finally, we examined if de-repression of *lin-41* was necessary for *let-7* mutant phenotypes, by asking whether restoring *let-7*-mediated silencing of *lin-41* but not of the other targets sufficed to suppress a given phenotype. To do so, we studied *lin-41(xe11); let-7(n2853)* double mutant animals (Ecsedi et al., 2015), in which both *let-7* target sites on the *lin-41* 3’UTR contain a compensatory point mutation to allow base pairing to the mutant version of *let-7* (Figure 1A). This results in substantial, albeit incomplete, repression of *lin-41*, while the other *let-7* target mRNAs are de-silenced to the same extent as in *let-7(PM)* single mutant animals (Figure 1A) (Aeschimann et al., 2017; Ecsedi et al., 2015). In the following, we refer to *lin-41(xe11)* as *lin-41(CPM)*, where *CPM* stands for compensatory point mutations.

**Figure 1.**
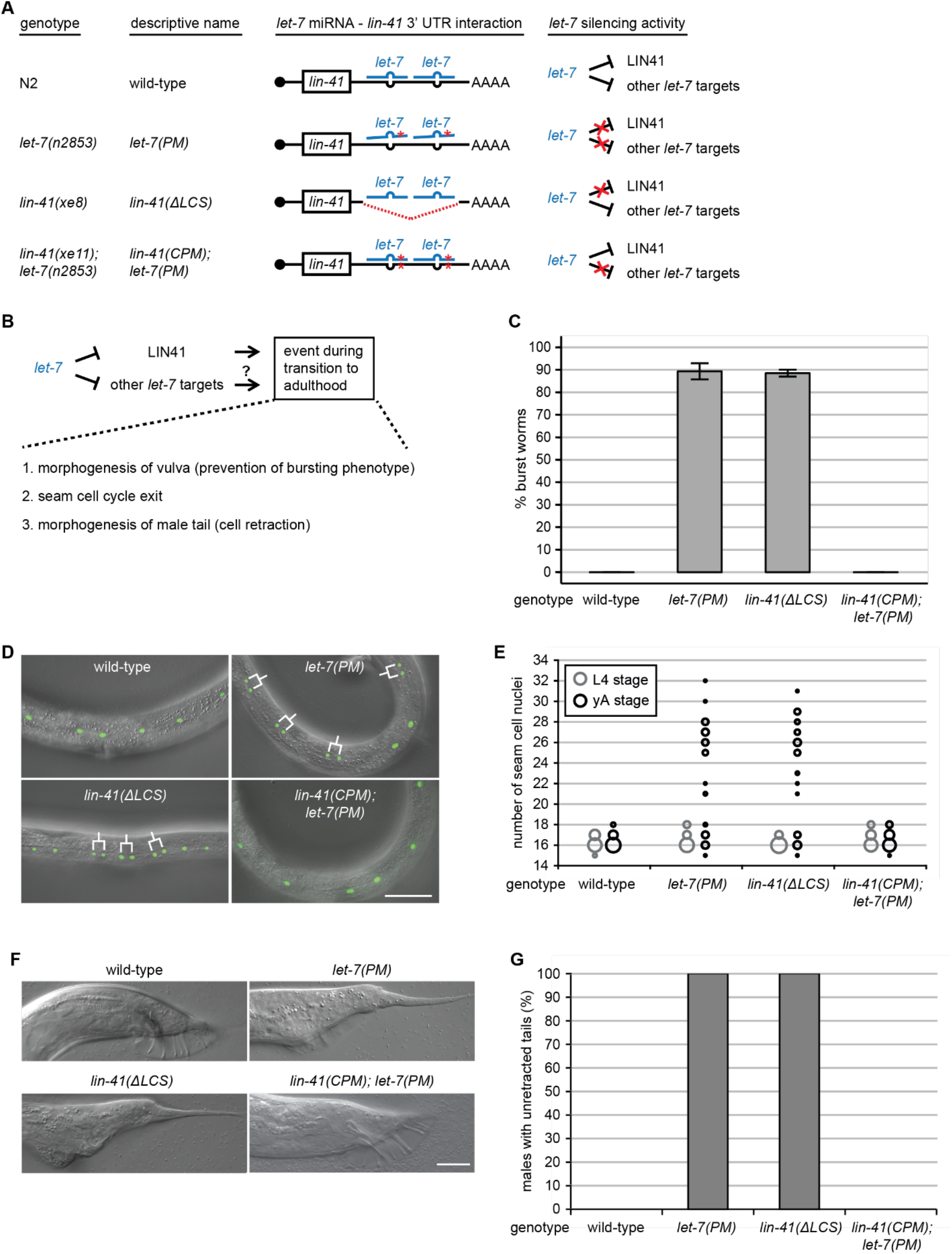
Failure of LIN41 downregulation explains multiple *let-7* phenotypes. (A) Schematic illustration of *let-7* and *lin-41* 3’UTR mutant alleles (not to scale) and of the *let-7* silencing activities in the different mutant backgrounds. Red asterisks depict point mutations and the red dotted line indicates a deletion. For both *let-7* target sites on the *lin-41* 3’UTR in the *lin-41(xe11); let-7(n2853)* background, a wild-type G:C base pair is replaced by an A:U base pair. This rescues *lin-41* downregulation by *let-7*, although not to the full extent (Aeschimann et al., 2017). (B) Schematic of a section of the heterochronic pathway regulating the onset of events during the larval-to-adult transition. The experiments of Figure 1 test if these events are regulated by silencing of only one *let-7* target (LIN41) or by silencing of any other combination of *let-7* targets. (C) The percentage of burst adult worms in synchronized populations grown at 25 °C for 45 hours of the indicated genotypes. Data as mean ± s.e.m of N = 3 independent biological replicates with n≥400 worms counted for each genotype and replicate. (D) Example micrographs of young adult worms expressing transgenic *scm::gfp* to visualize seam cell nuclei. Branched lines indicate seam cells originating from extra cell divisions. Scale bars: 50 μm. (E) Quantification of seam cell nuclei at the late L4 larval (L4) or young adult (yA) stage, in worms of the indicated genetic backgrounds. Areas of bubbles represent the percentage of worms with the corresponding number of seam cells. n=20 for L4, n>50 for yA, worms grown at 25 °C. (F) Example micrographs of tails in adult males of the indicated genetic background. Scale bars: 20 μm. (G) The percentage of young adult males of the indicated genotype with unretracted tails. n≥100, worms grown at 25 °C.

This set of genetic backgrounds allowed us to test the “one target hypothesis” for each developmental event *let-7* regulates (Figure 1B). Specifically, in addition to the role of *let-7* in vulva morphogenesis, we wanted to explore whether control of seam cell self-renewal and male tail retraction by *let-7* also solely depended on silencing of *lin-41*. Replicating our previous results (Ecsedi et al., 2015), the failure to downregulate *lin-41* explained the lethality of the *let-7* mutation, as this phenotype was recapitulated in *lin-41(∆LCS)* animals and rescued in *lin-41(CPM); let-7(PM)* animals (Figure 1C). We next examined *let-7*-mediated control of seam cell proliferation. Loss of *let-7* activity results in a failure of seam cells to exit the cell cycle (Reinhart et al., 2000), resulting in additional seam cell divisions at the young adult stage. We quantified this phenotype by counting the number of GFP-marked seam cell nuclei at the late L4 stage and a few hours later, in young adults shortly before *let-7(PM)* and *lin-41(∆LCS)* animals died. Both *lin-41(∆LCS)* and *let-7(PM)* mutant animals exhibited a wild-type number of seam cells at the L4 stage, just before the final molt (Figure 1E, gray circles). However, both mutations caused a failure in termination of the self-renewal program at the transition to adulthood, such that, after the molt, mutant animals had more seam cells than wild-type animals (Figure 1E, black circles; Table S1). As *lin-41(∆LCS)* recapitulated the *let-7(PM)* phenotype qualitatively and quantitatively (Figures 1D, 1E, Table S1), we conclude that dysregulation of *lin-41* is sufficient for this *let-7* mutant phenotype. Moreover, because restored repression of *lin-41* in *lin-41(CPM); let-7(PM)* double mutant animals sufficed to revert seam cell numbers to the lower, wild-type level (Figures 1D, 1E, Table S1), *lin-41* dysregulation is also necessary for the phenotype. Thus, LIN41 is the single target of *let-7* in controlling seam cell self-renewal.

Next, we analyzed the tails of adult males. Both *let-7(PM)* and *lin-41* gain-of-function (gf) males were reported to display partial defects in tail retraction (Del Rio-Albrechtsen et al., 2006). However, *let-7(PM)* mutants were analyzed only at 15°C, a semi-permissive temperature for survival and presumably other phenotypes. Moreover, the *lin-41gf* alleles used were also considered weak, and the mechanistic basis of their hyperactivation, involving mutations in the first coding exon of *lin-41*, is unknown. Strikingly, using *let-7(PM)* animals at 25°C and *lin-41(∆LCS)* mutant animals, we observed much more severe phenotypes. As analyzed in more detail in the following section, almost all *let-7(PM)* and all *lin-41(∆LCS)* mutant adult males exhibited “unretracted” tails with a long, pointed shape, resembling those of hermaphrodites (Figures 1F, 1G, Table S2). This phenotype was again caused by a failure of *let-7* to downregulate *lin-41*, as the *let-7(PM)* phenotype was suppressed in *lin-41(CPM); let-7(PM)* double mutant animals (Figures 1F, 1G, Table S2). We conclude that all three analyzed *let-7* mutant phenotypes can be fully explained by dysregulation of *lin-41*. Hence, *lin-41* is the key *let-7* target for different developmental functions and in different tissues.

### Sustained LIN41 expression leads to a complete failure of male tail cell retraction

We were struck by the severity of the mail tail retraction phenotype that we observed in *lin-41(∆LCS)* and *let-7(PM)* mutant animals and investigated this phenotype in more detail.

During the L4 stage, male tails are re-shaped from a pointed into a more rounded structure, as a consequence of cells moving anteriorly (Nguyen et al., 1999). In wild-type males, the first visible step occurs at the mid L4 stage, when the epidermal cell hyp8-11 at the very tip of the tail retracts anteriorly, withdrawing from the larval cuticle and turning the tip into a rounded shape (Figure 2A(ii), dashed arrow). At the late L4 stage, the entire tail retracts, leaving behind male-specific sensilla called rays (Figure 2A(iii), arrow). After the last molt, the adult male tail is freed from the larval cuticle, but keeps a structure called the fan (Figure 2A(iv), arrowhead), consisting of a fold in the outer layer of the adult cuticle.

**Figure 2.**
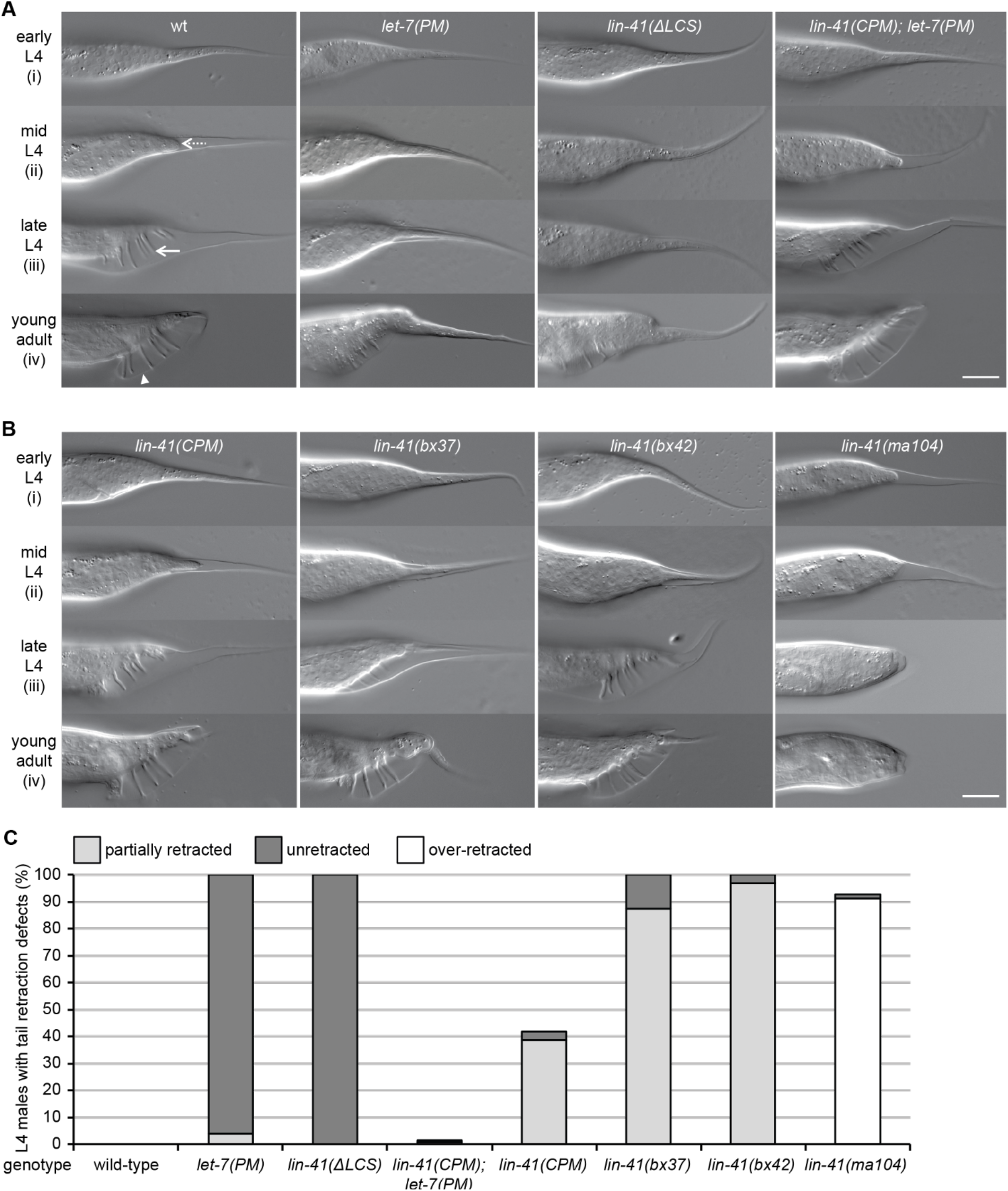
A failure in LIN41 downregulation results in a complete loss of male tail cell retractions. (A, B) Example micrographs of male tails at different developmental time points in the indicated genetic backgrounds. The dashed arrow illustrates the anterior retraction of the epidermal cell at the tail tip. The full arrow and the arrowhead point to one of the rays and the fan, respectively. Scale bars: 20 μm. (C) Quantification of the male tail phenotypes of the indicated genotypes at the late L4 larval stage as illustrated with pictures (iii) in (A, B). Shown are the percentages of animals with over-retracted, partially retracted or unretracted tails. n≥100, except for *lin-41(ma104)* (n=80). Worms were grown at 25 °C. (A-C) *lin-41(ma104)* animals display a precocious male tail retraction phenotype and were included as a control.

As previously reported (Del Rio-Albrechtsen et al., 2006), the weak gf mutations *lin-41(bx37)* and *lin-41(bx42)* caused a “leptoderan” (Lep) phenotype (Figure 2B(iv), Table S2). The Lep phenotype is characterized by a selectively delayed retraction of hyp8-11 but not of the other cells, giving rise to animals with tails that appear wild-type in shape except for the presence of a tail spike (Nguyen et al., 1999). By contrast, rather than being merely delayed, tail tip cell retraction failed entirely in both *lin-41(∆LCS)* and *let-7(PM)* males, (Figures 2A, 2C; Table S2). Thus, while most *lin-41(bx37gf)* or *lin-41(bx42gf)* mutant males had a partially retracted tail at the late L4 stage, with a spike protruding from the otherwise normal tail, none of the *lin-41(∆LCS)* mutant males displayed any cell retraction (Figures 2A(iii), 2B(iii), 2C, Table S2). At the young adult stage, this resulted in Lep tails for *lin-41(bx37)* or *lin-41(bx42)* mutants, whereas *lin-41(∆LCS)* and *let-7(PM)* animals had a distinct and more severe “unretracted” phenotype (Figures 2A(iv), 2B(iv), S1B; Table S2).

From this analysis, we conclude that male tail retraction absolutely depends on properly timed repression of *lin-41* by *let-7*. Normal execution of the program in *lin-41(CPM); let-7(PM)* double mutant animals confirms that the two genes function as a regulatory module also in this pathway (Figures 2A, 2C, Table S2). Finally, we note that male tail morphogenesis appears particularly sensitive to LIN41 activity levels, since the weak *gf* alleles *lin-41(bx37)* and *lin-41(bx42)* cause delayed, albeit largely functional male tail retraction (Figure 2B) but no seam cell division or survival phenotypes (Del Rio-Albrechtsen et al., 2006). The same is true for the *lin-41(CPM)* single mutant (Figure 2B), where *lin-41* is partially refractory to repression by wild-type *let-7*, causing a moderate *lin-41* de-silencing.

### LIN41 specifically binds to only a few somatic mRNAs

Since it was the failure to silence *lin-41* that caused *let-7* mutant phenotypes (Figure 1), we aimed at characterizing the relevant physiological roles of LIN41. To this end, we performed RNA co-immunoprecipitation coupled to RNA sequencing (RIP-seq) on a semi-synchronous population of animals in the L3 and L4 larval stages, during which LIN41 regulates the four previously identified mRNA targets. Using an anti-FLAG antibody, we applied RIP-seq to worms with transgenic expression of a rescuing *flag::gfp::lin-41* transgene in the *lin-41(n2914)* mutant background (Aeschimann et al., 2017) and to wild-type worms expressing *flag::gfp::sart-3* (Rüegger et al., 2015) as a control. To identify candidate targets for all three LIN41 functions that we studied, we performed RIP-seq on worm populations enriched for males, using the *him-5(e1490)* genetic background to achieve a ~35% frequency of males in the population, compared to < 1% in the wild-type background (Meneely et al., 2012).

We performed three independent experiments and determined a set of LIN41-bound mRNAs using edgeR (Robinson et al., 2010) with a false discovery rate of < 0.05. This set contained only seven mRNAs, the unannotated *F18C5.10* and *C31H5.5*, *lin-29a*, *mab-10*, *mab-3, dmd-3*, and *lin-41* (Figure 3A). The identification of *lin-41* mRNA in the IPs could reflect direct binding of LIN41 to its own mRNA to regulate it. Previously, LIN41 had been proposed to auto-regulate its activity (Del Rio-Albrechtsen et al., 2006), although post-translationally rather than on the transcript level. While it remains possible that LIN41 regulates its own expression at the level of mRNA silencing, we consider it likely that the detection of *lin-41* mRNA in the immunoprecipitates results from the immunoprecipitation of nascent FLAG::GFP::LIN41 protein, still bound to the translating ribosome and thus, indirectly, to its own mRNA.

**Figure 3.**
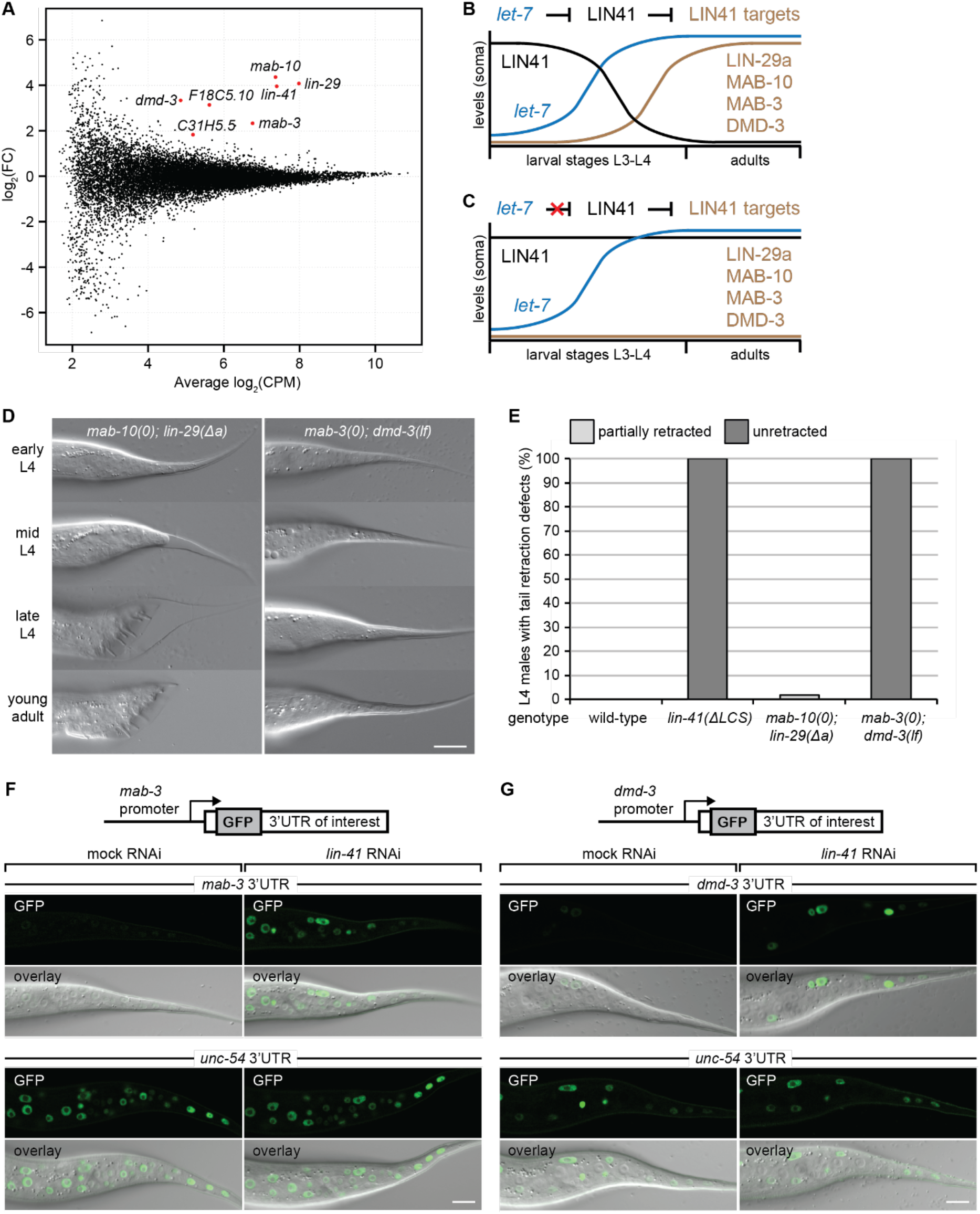
LIN41 controls male tail cell retraction through MAB-3 and DMD-3. (A) MA plot for anti-FLAG RNA co-immunoprecipitation coupled to RNA sequencing (RIP-seq) experiments with FLAG::GFP::LIN41 and FLAG::GFP::SART-3 as a control. Semi-synchronous L3/L4 stage worm populations enriched in males (*him-5(e1490)* genetic background) were compared in three independent biological replicates. The plot compares the fold change (FC) in IP-to-input enrichments for RNA-sequencing reads in the FLAG::GFP::LIN41 versus the FLAG::GFP::SART-3 IP (y-axis) with the mean mRNA abundance (x-axis, CPM: counts per million). Genes passing the cutoff of FDR < 0.05 are highlighted in red and labeled. (B, C) Schematic expression of the LIN41 protein, the *let-7* miRNA, and the LIN41 target proteins in the soma during development from larvae to adult worms, in the wild-type situation (B) and when *let-7* fails to repress *lin-41* (C). (D) Example micrographs of male tails at different developmental time points in the indicated genetic backgrounds. Scale bars: 20 μm. (E) Quantification of the male tail phenotypes at the late L4 larval stage of *mab-10(0) lin-29(Δa)* and *mab-3(0); dmd-3(lf)* animals. The data for wild-type and *lin-41(∆LCS)* males are re-plotted from Figure 1 for reference. Shown are the percentages of animals with over-retracted, partially retracted or unretracted tails. n≥100, worms were grown at 25 °C. (F, G) Confocal images of the male tail epidermis in young L3 stage male animals expressing nuclear-localized GFP(PEST)::H2B reporters, driven from the *mab-3* (F) and *dmd-3* (G) promoters and fused to their orthologous 3’UTR sequences or the unregulated *unc-54* 3’UTR as indicated. Animals were grown for 20 h at 25 °C on *lin-41* or mock RNAi bacteria. Scale bars: 10 μm.

Strikingly, the remaining six mRNAs included *lin-29a*, *mab-10*, *mab-3* and *dmd-3*. These were the only four mRNAs that we had previously characterized as LIN41 targets on the molecular level after ribosome profiling experiments had revealed their repression in mutant animals with sustained LIN41 expression (Aeschimann et al., 2017). Because *F18C5.10* and *C31H5.5* were unchanged in these same ribosome profiling experiments, we examined whether regulation of *lin-29*, *mab-10*, *mab-3* and/or *dmd-3* mRNAs could explain the physiological functions of LIN41.

LIN-29 is an early growth response (EGR)-type transcription factor of the *Krüppel* family and occurs in two major isoforms, LIN-29a and LIN-29b (Rougvie and Ambros, 1995). MAB-10, orthologous to mammalian NAB1/2, is a co-factor of LIN-29 and can associate with both isoforms (Harris and Horvitz, 2011). MAB-3 and DMD-3 are both DM (<u>D</u>oublesex/<u>M</u>AB-3) domain-containing transcription factors, proposed to act at least partially redundantly on common targets due to similar binding motifs (Mason et al., 2008; Yi and Zarkower, 1999). Therefore, we wondered if the main function of the *let-7 –* LIN41 module was to control the activity of two pairs of transcriptional regulators; i.e., LIN-29 + MAB-10 and MAB-3 + DMD-3, respectively. If this were true, the changes in activity of these LIN41 targets should explain the phenotypes we observed in *let-7(PM)* and *lin-41(∆LCS)* mutant animals.

Previously, we showed that all four transcriptional regulators accumulate shortly before wild-type worms turn into adults, when *let-7* has silenced LIN41 sufficiently to prevent repression of LIN41 targets ((Aeschimann et al., 2017) & schematically depicted in Figure 3B). Accordingly, when *let-7*-mediated silencing of LIN41 is lost, e.g., in *let-7(PM)* and *lin-41(∆LCS)* mutant animals, sustained LIN41 accumulation keeps the four LIN41 targets repressed (Figure 3C). We therefore asked whether the failure to express one or several of these targets could explain the phenotypes of *let-7(PM)* and *lin-41(∆LCS)* animals; i.e., whether we could recapitulate these phenotypes by inactivating the transcriptional regulators genetically.

### LIN41 regulates MAB-3 and DMD-3 to time male tail retraction

To explore the effects of losing the activity of LIN41 targets, we used the previously described *dmd-3(ok1327)* allele, referred to as *dmd-3(lf)* but considered null for its function in male tail development (Mason et al., 2008), and generated null alleles for the other three genes using CRISPR-Cas9. Specifically, for *mab-10* and *mab-3*, we deleted almost the entire coding region including all functionally important domains (Figure S1A). For *lin-29*, we previously established that LIN41 targets only one *lin-29* isoform, *lin-29a*, without targeting the *lin-29b* isoform (Aeschimann et al., 2017). Hence, we specifically disrupted *lin-29a*, without affecting expression of *lin-29b*. The resulting allele deletes coding exons two through four of *lin-29* while causing a frame-shift in the remaining *lin-29a* open reading frame, rendering it null for *lin-29a* (Figure S1A).

First, we examined male tail retraction. Mating deficiencies and abnormalities in the tail tips have been reported for males mutant for each of these four LIN41 targets, although cell retraction appears to occur normally in *lin-29* and *mab-10* single mutant males (Euling et al., 1999; Hodgkin, 1983). However, *mab-3* and *dmd-3* single mutant males exhibit Lep phenotypes, which are weak and of low penetrance for *mab-3* but stronger and highly penetrant for *dmd-3* (Mason et al., 2008). Moreover, *mab-3; dmd-3* double mutant males have completely unretracted male tails (Mason et al., 2008). In order to compare phenotypes of *lin-41(∆LCS)* mutants to those of the LIN41 target pairs, we observed and quantified the male tail phenotypes of *mab-10(0) lin-29(Δa)* as well as of *mab-3(0); dmd-3(lf)* double mutant animals. While the *mab-10(0) lin-29(Δa)* males had short fan and ray structures, these animals were wild-type for tail cell retraction (Figures 3D, 3E, S1B, Table S2). By contrast and confirming earlier results (Mason et al., 2008), *mab-3(0); dmd-3(lf)* males showed no sign of cell retraction at any stage during development (Figures 3D, 3E, S1B, Table S2). This phenocopies the *lin-41(∆LCS)* and *let-7(PM)* mutations (Figures 1F, 2A, 2C, Table S2).

Previously, it was proposed that *dmd-3* expression was indirectly regulated by LIN41, through an unknown mechanism involving the *dmd-3* promoter (Mason et al., 2008). By contrast, using ectopic reporter gene expression in the hermaphrodite epidermis, we found that both *mab-3* and *dmd-3* 3’UTRs confer LIN41-dependent regulation (Aeschimann et al., 2017). To test whether physiologic regulation of *mab-3* and *dmd-3* by LIN41 in the male tail epidermis was promoter-dependent or direct and 3’UTR-dependent, we created reporter lines to express *gfp(pest)::h2b* from putative *mab-3* and *dmd-3* promoters, i.e. the 4 kb region upstream of their start codons. For each promoter, we created two reporters, one with a promoter-orthologous 3’UTR that harbors LIN41 target sites and one with the heterologous, unregulated *unc-54* 3’UTR (Aeschimann et al., 2017).

Whereas all four reporters were expressed in the tail epidermis and other tissues of males at the L4 stage, when LIN41 levels are low due to the down-regulation by *let-7*, differences occurred in early L3 stage males, when LIN41 levels are high: GFP expression in most tissues, and in particular in the epidermis, was almost undetectable for both reporters carrying the promoter-orthologous 3’UTRs (Figures 3F, 3G). By contrast, both reporters containing the heterologous *unc-54* 3’UTR revealed strong GFP expression in various cells, including the epidermal cells of the tail region (Figures 3F, 3G & data not shown). Moreover, for both reporters with the promoter-orthologous 3’UTRs, depletion of LIN41 by RNAi resulted in strong GFP signals in many tissues, including the epidermal cells of the tail region (Figures 3F, 3G). We conclude that temporal control of MAB-3 and DMD-3 accumulation is predominantly conferred by posttranscriptional regulation, through LIN41-binding to their 3’UTRs and, hence, that LIN41 regulates the timing of cell retraction in the male tail through *mab-3* and *dmd-3* mRNAs as direct targets.

### Seam cell exit from the cell cycle is controlled by LIN41-mediated regulation of LIN-29a and MAB-10

Since sustained LIN41 expression leads to a failure in cell cycle exit of seam cells, we next examined the functions of the LIN41 targets in this process. Both *lin-29* and *mab-10* have been implicated in seam cell development (Ambros and Horvitz, 1984; Harris and Horvitz, 2011), so we wondered if we could phenocopy *let-7(PM)* or *lin-41(∆LCS)* mutants by mutating *lin-29a* and/or *mab-10*. Seam cell numbers in young adults of either *lin-29(Δa)* or *mab-10(0)* single mutants were unchanged from the wild-type situation (Figure 4A, Table S1), although older *mab-10(0)* adults displayed a few additional seam cells (data not shown & (Harris and Horvitz, 2011)). By contrast, young adult *mab-10(0) lin-29(Δa)* double mutant animals displayed additional seam cell divisions to a comparable extent as *let-7(PM)* or *lin-41(∆LCS)* mutant animals (Figures 1D, 4A, 4B, Table S1). Depletion of both MAB-3 and DMD-3 did not affect seam cell numbers (Figure 4A, Table S1). We conclude that *lin-29a* and *mab-10* are a major, likely sole, output of the *let-7–*LIN41 regulatory module for control of seam cell self-renewal. We also note that the synthetic *mab-10(0) lin-29(∆a)* mutant phenotype indicates that LIN-29a cannot be the only transcription factor the activity of which MAB-10 modulates. For cessation of seam cell self-renewal, the relevant additional transcription factor is most likely LIN-29b. This is because *lin-29(0)* mutant animals, lacking both LIN-29 isoforms but not MAB-10, exhibit a penetrant defect in cessation of seam cell self-renewal (Table S1), and because MAB-10 can bind to both LIN-29 isoforms. Hence, it appears that LIN41 promotes seam cell self-renewal through combining direct repression of LIN-29a with indirect control over the activity of both LIN-29a and LIN-29b by repression of their shared co-factor MAB-10.

**Figure 4.**
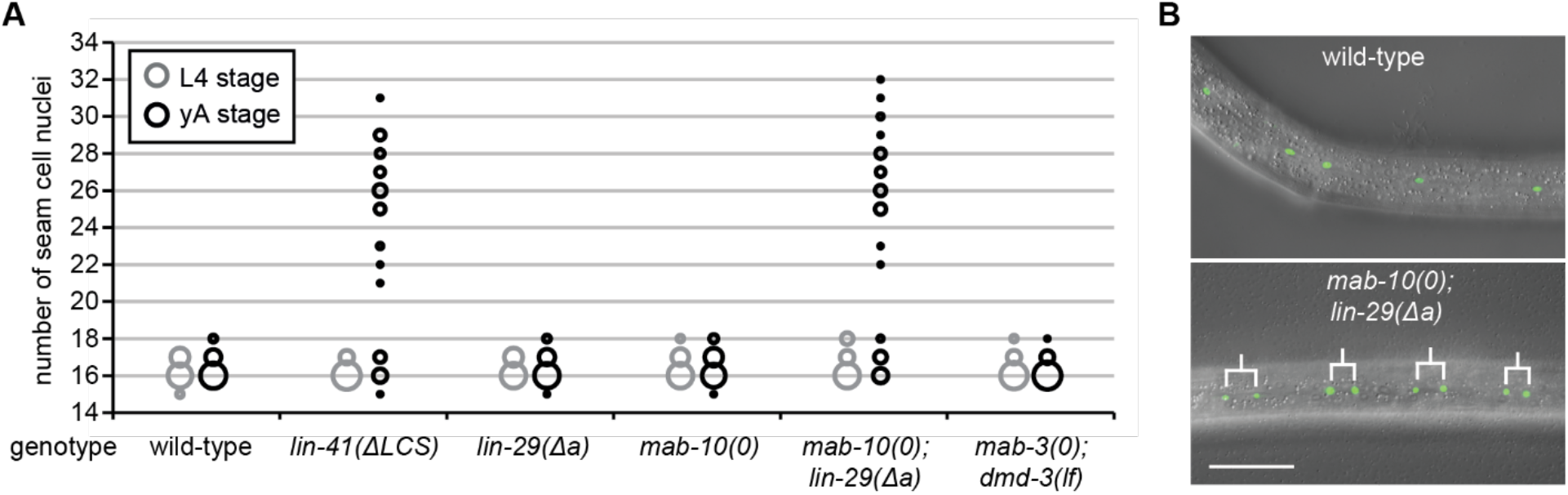
LIN41 controls self-renewal of seam cells through LIN-29a and its co-factor MAB-10. (A) L4 larval stage and young adult (yA) worm seam cell nuclei quantifications in indicated genetic backgrounds, as in Figure 1D. n=20 for L4, n>50 for yA, worms grown at 25 °C. Results for wild-type and *lin-41(∆LCS)* animals are re-plotted from Figure 1 for reference. (B) Example micrographs of a young adult wild-type worm and a worm depleted of LIN-29a and MAB-10 expressing transgenic *scm::gfp*. Branched lines indicate seam cell nuclei originating from extra cell divisions. Scale bar: 50 μm.

### Depletion of LIN-29a and MAB-10 causes vulval bursting

Whereas elevated LIN41 levels can fully explain why *let-7* mutant worms are inviable ((Ecsedi et al., 2015) and Figure 1B), it is unknown why these elevated LIN41 levels cause hermaphrodites to burst through the vulva. Since the *Drosophila* LIN41 orthologue dappled/wech links integrins to the cytoskeleton (Loer et al., 2008), LIN41 might directly contribute to the structural integrity of the vulva, rather than promote its morphogenesis through post-transcriptional gene regulation (Ecsedi et al., 2015). Consistent with this idea, vulval bursting has not been reported for mutants of *lin-29*, *mab-10*, *mab-3* or *dmd-3*. Nevertheless, we considered the possibility that isoform-specific dys-regulation of LIN-29a, or a combination of dys-regulated LIN41 targets would cause vulval rupturing shortly after the larval-to-adult transition.

When we examined *mab-3(0)*; *dmd-3(lf)* double mutant animals, we observed no bursting (Figure 5A, Table S3). The same was true for *mab-10(0)* single mutant animals. However, isoform-specific deletion of LIN-29a resulted in occasional (< 2 %) bursting, and the penetrance of this phenotype increased to 15 % in *mab-10(0) lin-29(∆a)* double mutant animals (Figure 5A, Table S3). Hence, a combined dysregulation of LIN41 targets appeared to account for the bursting phenotype. Yet, surprisingly, the penetrance of vulval bursting was not further increased even when we examined *mab-3 mab-10 lin-29(∆a); dmd-3* quadruple mutant animals (Figure 5A, Table S3).

**Figure 5.**
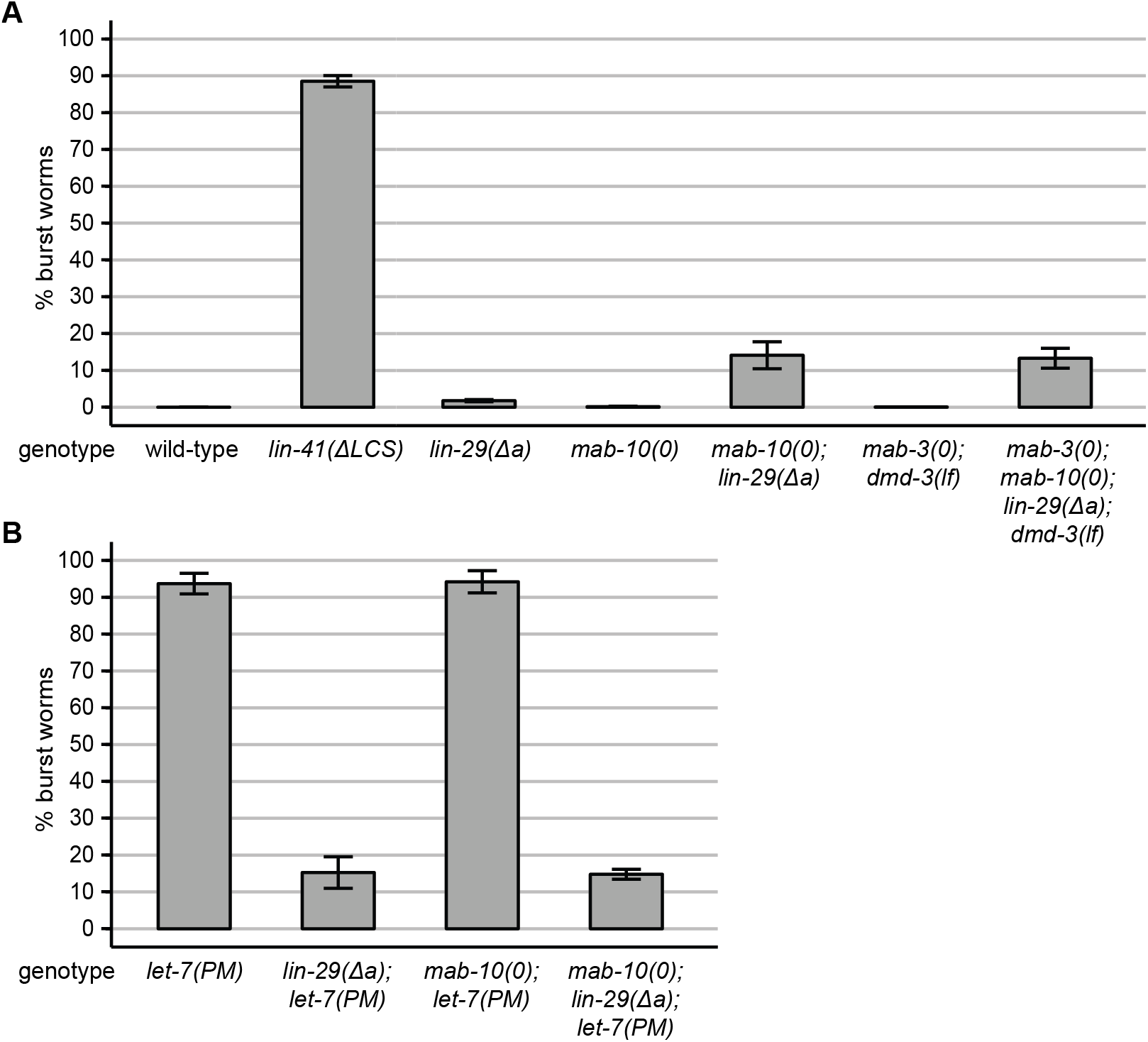
LIN-29a and MAB-10 are involved in the vulva bursting phenotype. (A, B) Quantification of burst worms of the indicated genotypes, grown in a synchronized population at 25 °C for 45 hours. Data as mean ± s.e.m of N = 3 independent biological replicates with n≥400 worms counted for each genotype and replicate. In panel (A), results for wild-type and *lin-41(∆LCS)* animals are re-plotted from Figure 1 for reference.

Conceivably, failure of the quadruple mutant to fully recapitulate the *let-7(PM)* and *lin-41(∆LCS)* bursting phenotypes, which are ~90 % penetrant at the same developmental stage, might reflect the contribution of one or several more LIN41 targets that we have thus far failed to identify, or a structural role of the LIN41 protein. However, we considered this unlikely because we observed that *mab-10(0) lin-29(∆a); let-7(PM)* triple mutant animals phenocopied the *mab-10(0) lin-29(∆a)* double mutant rather than the *let-7(PM)* single mutant phenotype (Figure 5B). In other words, mutations in *mab-10* and *lin-29a* suppressed the *let-7(PM)* mutant phenotype. The suppression was due to the *lin-29(Δa)* and not the *mab-10(0)* mutation, as *lin-29(Δa)* alone reduced the bursting frequency of *let-7(PM)* animals to about 15 %, while *mab-10(0)* did not reduce the bursting frequency (Figure 5B, Table S3). Hence, it appeared that complete inactivation of *lin-29a* through genetic deletion interferes with recapitulation of the *let-7(PM)* mutant phenotype, and that a residual or tissue-specific activity might be necessary to permit vulval bursting with high penetrance.

### Lin-29a is expressed in the anchor cell of wild-type and let-7 mutant animals

In order to observe when and where LIN-29a is differentially expressed in wild-type and *let-7(PM)* worms, we specifically tagged the endogenous LIN-29a isoform by placing a GFP::3xFLAG tag at the N-terminus (Figure S1A). As expected, we observed *lin-29a* expression in the epidermis of wild-type but not *let-7(PM)* mutant late L4 stage animals (Figure 6A). However, not all *lin-29a* expression was lost in the *let-7(PM)* mutant. Specifically, we observed GFP::LIN-29a accumulation in the distal tip cell, the anchor cell (AC) and most vulval cells. The strongest GFP signal was that in the AC (Figure 6B), observed from about the L2-to-L3 molt onwards.

**Figure 6.**
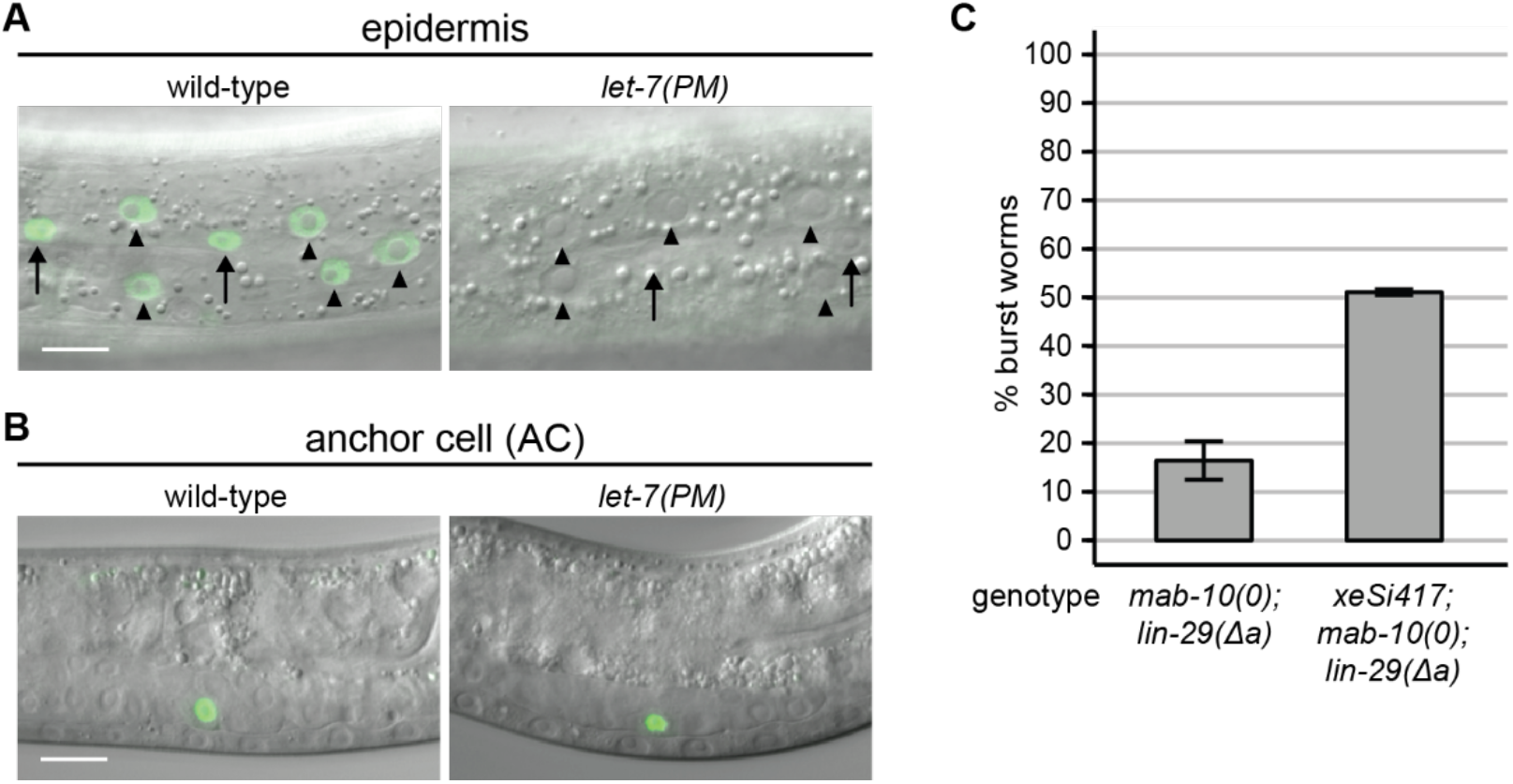
LIN-29a accumulation in the AC is independent of *let-7*. (A, B) Example micrographs of worms expressing GFP-tagged LIN-29a, in the epidermis of late L4 stage animals (A) and in the anchor cell of L3 stage animals (B). Scale bars: 10 μm. Arrows point to seam cell nuclei, arrowheads to hyp7 nuclei. (C) Quantification of burst worms of the indicated genotypes, grown in a synchronized population at 25 °C for 45 hours. The xeSi417 transgene drives *lin-29a* expression specifically in the anchor cell. Data as mean ± s.e.m of N = 3 independent biological replicates with n≥400 worms counted for each genotype and replicate.

We considered expression in the AC particularly interesting, given that the AC establishes the uterine-vulval connection through which *let-7* mutant animals burst. Moreover, and as discussed in more detail in the following section, *lin-29* mutations cause defects in vulval-uterine cell fates that are known to be AC-dependent (Newman et al., 2000), and that are not shared by *let-7* mutant animals (Ecsedi et al., 2015). Therefore, we wondered if LIN-29a activity in the AC contributed to the increased bursting frequency of *let-7(PM)* relative to *mab-10(0) lin-29(∆a)* mutant animals. To investigate this, we re-expressed LIN-29a specifically in the AC of *mab-10(0) lin-29(∆a)* mutant animals by combining the *∆pes-10* basal promoter (Harfe and Fire, 1998) with an anchor cell-specific enhancer of *lin-3* (ACEL, (Hwang and Sternberg, 2004)). The single-copy integrated *xeSi417* transgene thus generated encodes an operon containing *lin-29a* followed by nuclear-localized *gfp* to assay the expression pattern. We observed consistent and specific anchor cell expression, although the GFP levels failed to reach those of the endogenously tagged GFP::LIN-29a protein. Wild-type animals expressing the transgene did not reveal any overt defects. However, consistent with our hypothesis, expression of this transgene in *mab-10(0) lin-29(Δa)* mutant animals increased the frequency of bursting to about 50 % (Figure 6C). We conclude that LIN-29a expression in the AC of animals otherwise lacking LIN-29a and its co-factor MAB-10 are more prone to vulval rupturing than those lacking LIN-29a and MAB-10 in all tissues. Although the experiment did not result in complete penetrance of the bursting phenotype, we further conclude that *let-7(PM)* mutant animals likely die due to a lack of LIN-29 activity in some tissues while retaining LIN-29 activity in the anchor cell.

### Lin-29a expression in the AC promotes uterine seam cell formation

Given the striking increase in vulval bursting of *mab-10(0) lin-29(∆a)* mutant animals when they expressed *lin-29a* in the AC, we sought to examine the role of LIN-29a in the AC further. During wild-type development, the AC induces and coordinates cell fates of both the vulva and the uterus. In uterine development, the AC specifies its lateral neighbors as π cells, and some of the π daughter cells fuse with each other and eventually with the AC to form the uterine seam cell (utse). The utse is important for the structure of the egg-laying apparatus, as it anchors the adult uterus to the epidermal seam. It also forms a thin cytoplasmic extension that separates the lumens of vulva and uterus until it is broken when the first embryo is laid. In *lin-29(0)* mutants, π cell fates are not specified and the utse does not form. Instead, the AC remains unfused, and the vulva and uterus are separated by thick tissue rather than the thin utse (Newman et al., 2000); Figure 7A). When *let-7* mutant animals die by bursting through the vulva, the intestine is pushed out of the animal, which breaks the thin utse. We hypothesized that lack of *lin-29a* expression in the AC of *mab-10(0) lin-29(Δa)* mutant animals could also lead to a failure in utse formation, and that a thicker tissue could prevent vulval bursting. Indeed, we found that more than 50 % of *mab-10(0) lin-29(Δa)* mutant hermaphrodites had a thick tissue separating the vulval and uterine lumens. Moreover, re-expressing *lin-29a* in the AC in *mab-10(0) lin-29(Δa); xeSi417* hermaphrodites restored utse formation in 100% of animals as scored by re-appearance of the thin cytoplasmic extension (Figures 7A, 7B, Table S4). We conclude that expression of *lin-29a* in the AC contributes to both wild-type utse formation and vulval bursting.

**Figure 7.**
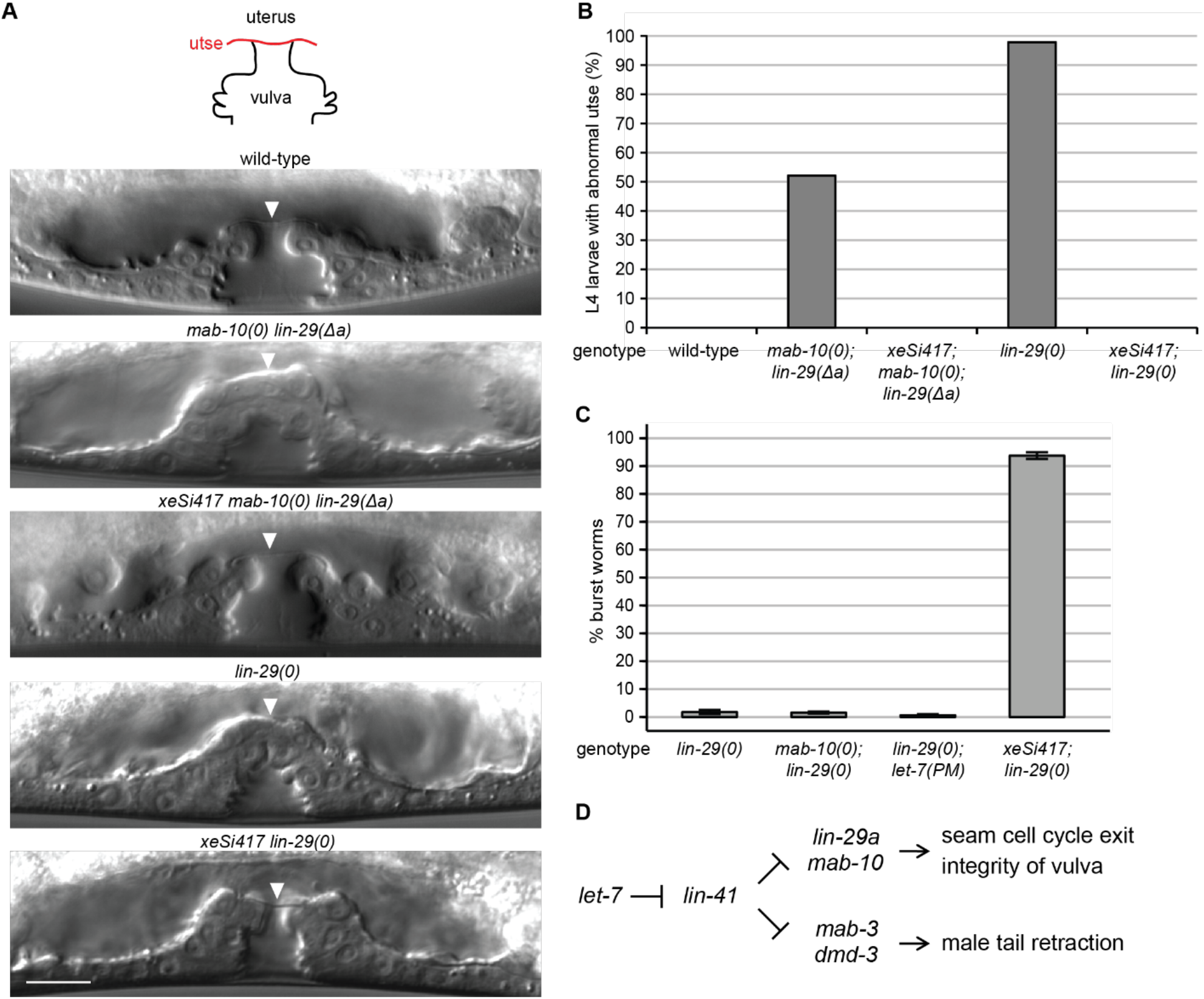
LIN-29a accumulation in the AC is required for utse formation. (A) Example pictures of the late L4 stage vulva in wild-type animals and *mab-10(0) lin-29(Δa)* or *lin-29(0)* mutant animals with and without expression of *lin-29a* in the anchor cell through the *xeSi417* transgene. Arrowheads point to the thin or thick tissue separating the vulva from the uterus. Scale bar: 10 μm. (B) Quantification of uterine-vulva connection phenotypes of late L4 larvae of the indicated genotypes. n≥40, worms grown at 25 °C. (C) Quantification of burst worms of the indicated genotypes, grown in a synchronized population at 25 °C for 45 hours. Data as mean ± s.e.m of N = 3 independent biological replicates with n≥400 worms counted for each genotype and replicate. (D) Model for the output of the *let-7* - LIN41 pathway regulating two pairs of LIN41 targets involved in transcription. LIN-29a and MAB-10 stop seam cell divisions and allow survival because vulval rupturing is prevented. MAB-3 and DMD-3 drive cell retraction events to shape the male tail.

Of the two LIN-29 isoforms, LIN41 regulates only LIN-29a. However, because it also regulates MAB-10, a common co-factor of LIN-29a and LIN-29b, one might expect that *lin-29(0)* mutations, affecting both isoforms, would cause phenotypes comparable to, or even more severe than, those of *let-7(PM)* and *mab-10(0) lin-29(∆a)* mutant animals. Consistent with this view, complete lack of LIN-29 activity in *lin-29(0)* mutant animals is well known to result in continued seam cell division at the adult stage (Table S1 & (Ambros and Horvitz, 1984)). However, *lin-29(0)* mutant animals have not been reported to exhibit vulval rupturing. To test this explicitly, we deleted almost the entire *lin-29* locus to generate the *lin-29(xe37)* mutant allele (Figure S1), subsequently referred to as *lin-29(0)*. We observed that *lin-29(0)* mutants rarely burst (Figure 7C). In fact, at < 2 %, the penetrance was similar to that of *lin-29(Δa)* mutant animals. However, contrasting with *lin-29(∆a)*, it was not further enhanced by loss of *mab-10* (Figure 7C). Hence, the more penetrant bursting phenotype of *mab-10(0) lin-29(∆a)* double mutant relative to *lin-29(0)* single mutant animals does not reflect an additional, LIN-29-independent function of MAB-10. Rather, it suggests that a complete loss of LIN-29 activity prevents vulval bursting. We confirmed this by crossing the *lin-29(0)* allele into *let-7(PM)* mutant animals, which suppressed the bursting frequency to < 2 % (Figure 7C).

We suspected that a utse formation defect, highly penetrant in animals lacking LIN-29 (Newman et al., 2000), prevented bursting of *lin-29(0)* mutant animals, as it did on *mab-10(0) lin-29(∆a)* double mutant animals. Indeed, re-expression of LIN-29a in the AC of *lin-29(0)* mutant animals restored wild-type utse morphology (Figures 7A, 7B, Table S4) and resulted in highly penetrant vulval bursting (Figure 7C). We conclude that a LIN-29a activity in only one cell, the AC, promotes utse formation, and that a complete lack of LIN-29 activity in all tissues but the AC is lethal to the worm.

## DISCUSSION

The *C. elegans* heterochronic pathway is arguably one of the best-characterized developmental timing pathways, with many of its genetic players and their relative positions in the pathway known. However, a limited knowledge of direct molecular connections among individual heterochronic genes has hampered a mechanistic understanding of this developmental timer. Building on previous knowledge that the RNA-binding protein LIN28 inhibits processing of the miRNA *let-7*, and *let-7* in turn silences target mRNAs post-transcriptionally (Rougvie and Moss, 2013), we have focused here on elucidating the molecular mechanisms that time transition into adulthood by identifying and functionally characterizing the players immediately downstream of *let-7*.

Extending our previous finding that *let-7* prevents vulval rupturing through regulation of only a single target (Ecsedi et al., 2015), LIN41, we find that its function in additional tissues and processes also relies on regulation of only LIN41. This finding is consistent with a previous gene expression analysis, which revealed that global, animal-wide changes in gene expression through *let-7* inactivation are recapitulated well by impairing selectively its silencing of only *lin-41* (Aeschimann et al., 2017). Yet, it contrasts with previous reports that identified numerous putative *let-7* targets (Abrahante et al., 2003; Andachi, 2008; Großhans et al., 2005; Hunter et al., 2013; Jovanovic et al., 2010; Lin et al., 2003). However, those reports relied on circumstantial evidence to establish miRNA targets, typically suppression of certain *let-7* mutant phenotypes through mutation or depletion of suspected target genes. The advent of genome editing has now enabled a direct analysis of the physiological relevance of an individual target, by specifically uncoupling it from *let-7*, or recoupling it to a mutant variant of *let-7*. The finding that the three *let-7*-regulated processes that we investigated – vulval rupturing, male tail morphogenesis and seam cell fate control – are all dependent on LIN41 as the key *let-7* target, now establishes *let-7* – LIN41 as a versatile regulatory module. Although miRNAs are often thought to act through a network activity where they silence many targets modestly but coordinately (Bracken et al., 2016), other instances have been reported where only one or two targets appear to explain physiological functions of miRNAs (Drexel et al., 2016; Lee et al., 1993; Moss et al., 1997; Wightman et al., 1993). Hence, direct testing of the network activity model seems warranted to understand whether it really reflects the predominant mode of miRNA function in animals.

The levels of LIN41 determine if and when certain developmental events are triggered. In mutants with too little or no LIN41 activity, events are triggered too early, and in mutants where LIN41 activity is too high, events are triggered too late or not at all. For instance, cell retraction at the male tail is premature with reduced-function alleles such as *lin-41(ma104)*, but it is delayed with gain-of-function mutations such as *lin-41(bx37)* or *lin-41(bx42)* (Figure 2 and (Del Rio-Albrechtsen et al., 2006)). Yet, previous analyses of *lin-41gf* animals or those expressing, presumably at elevated levels, *lin-41* from multicopy transgene arrays yielded only partially penetrant phenotypes for the various *let-7*-regulated processes (Del Rio-Albrechtsen et al., 2006; Slack et al., 2000). Hence, it was proposed that redundant *let-7* outputs existed (Slack et al., 2000). However, we would expect that *lin-41* activity in the previously examined overexpression and gf mutant strains remained under at least partial control of *let-7*. Accumulation of *let-7* would thus counteract LIN41 activity in late larval stages, although residual LIN41 levels might remain higher than in wild-type animals, or take longer to decline to a comparable level. In support of this view, tail retraction is only delayed when *let-7*-mediated silencing is reduced, but not entirely abrogated, in *lin-41(CPM)* males (Figure 2B) or in *let-7(PM)* males at the semi-permissive temperature of 15°C (Del Rio-Albrechtsen et al., 2006). By contrast, tail retraction is fully prevented when *lin-41* is entirely uncoupled from *let-7* by growing *let-7(PM)* animals at the restrictive temperature or by deleting its binding sites on the *lin-41* 3’UTR. Indeed, the elimination of temporal regulation of *lin-41* by *let-7* suffices to faithfully recapitulate the effect of complete *let-7* loss not only for male tail retraction but also for seam cell divisions and lethality.

Downstream of LIN41 in the heterochronic pathway, our results define the immediate next layer of regulatory function by identifying the relevant direct targets of LIN41 and their specific developmental roles. We demonstrate that *lin-29a*, *mab-10*, *mab-3* and *dmd-3*, previously shown to be directly, post-transcriptionally silenced by LIN41 (Aeschimann et al., 2017), act as the main regulatory output of the LIN28 – *let-7* – LIN41 pathway in the larval-to-adult transition. Thus, despite other proposed molecular activities, the function of LIN41 as an RNA-binding protein accounts for its known heterochronic functions. Moreover, in contrast to its rather promiscuous RNA-binding activity in the adult germline (Kumari et al., 2018; Tsukamoto et al., 2017) where its levels are high (Tocchini et al., 2014), LIN41 binding to mRNAs appears much more selective in the larval soma. Whether this is a consequence of the differences in protein concentration, as LIN41 accumulates to much higher levels in adult gonads than in larval soma, or of differential association with additional factors that modulate binding specificity in one situation or the other remains an open question.

Whereas *let-7* – LIN41 act sequentially, in a linear fashion, to control transition to adulthood, the heterochronic pathway branches at the point of LIN41 output (Figure 7D). This architecture facilitates a coordinated and timely activation of different developmental events as LIN41 becomes silenced through increasing *let-7* levels in late stage larvae. The LIN41 targets appear to be grouped into two functionally separated parallel pathways, each with a distinct pair of transcriptional regulators: DMD-3 + MAB-3 mediate male tail retraction, and LIN-29a + MAB-10 promote both vulval integrity and cessation of seam cell self-renewal. The individual transcription factors and co-factors within each pair seem to have partially redundant functions, as individual mutations cause either no, or only partially penetrant phenotypes. By co-regulating partially redundant genes within each pathway, LIN41 itself assumes a more unique function and, as a consequence, elevated LIN41 levels as in *lin-41(∆LCS)* mutants lead to fully penetrant phenotypes. Thus, to control events for the transition to adulthood, a cascade of post-transcriptional regulators eventually times the expression of two pairs of transcriptional regulators.

Although mating deficiencies and abnormalities in the male tail tips have been reported for individual mutant males of all four LIN41 targets (Del Rio-Albrechtsen et al., 2006; Euling et al., 1999; Hodgkin, 1983; Mason et al., 2008), cell retraction occurs normally *mab-10(0) lin-29(∆a)* double mutant animals. By contrast, male tail retraction is impaired in *dmd-3* single mutant animals and fails in *mab-3; dmd-3* double mutant animals (Mason et al., 2008). LIN41 had already previously been shown to regulate *dmd-3* expression, but regulation was thought to occur transcriptionally and indirectly, through the function of an unknown direct target of LIN41 (Mason et al., 2008). However, we recently showed that LIN41 can regulate *dmd-3* directly and post-transcriptionally, and that it also regulates *mab-3* in a similar manner (Aeschimann et al., 2017). This not only suggested a different mode of regulation, but also seemed incongruent with the much less severe phenotype previously reported for *lin-41gf* and *let-7lf* mutant animals compared to the *mab-3; dmd-3* double mutant animals (Del Rio-Albrechtsen et al., 2006; Mason et al., 2008). We can now reconcile these results by showing that *let-7 –* LIN41 are absolutely essential for male tail retraction, consistent with post-transcriptional regulation of both *dmd-3* and *mab-3* by LIN41.

Among the three mutant phenotypes that we have investigated, vulval rupturing is the most enigmatic. Although the phenotype was first described nearly two decades ago (Reinhart et al., 2000), its cause has remained unclear. As we now show, combined dysregulation of *lin-29a* and *mab-10* is an important factor. Since *mab-10(0) lin-29(∆a)* double mutant animals did not exhibit a fully penetrant phenotype, we cannot exclude that mis-expression of additional, yet to be discovered LIN41 targets, or a potential role of LIN41 as a structural protein might additionally contribute. However, based on the analysis of *lin-29a* expression and function in the anchor cell, we favor a different scenario, where sustained *lin-29a* expression in the anchor cell combines with the absence of *lin-29a* and *mab-10* in other, yet to be determined cells of the vulval-uterine system and/or the epidermis to cause penetrant vulval bursting.

Loss of *lin-29a* activity in the AC leads to defects in π-cell differentiation and thus failure of utse formation. The thick cell layer that instead separates uterus and vulva in mutant animals is presumably more resistant than a wild-type utse to the internal pressure in the worm, thus helping to avoid vulval bursting in animals that globally lack *lin-29a* and *mab-10* or both isoforms of *lin-29*. However, generation of a thin utse appears insufficient to permit bursting as re-expression of *gfp::lin-29a* in the anchor cell of *mab-10 lin-29a* increased vulval bursting to 50 % of animals while restoring overtly normal utse morphology to all animals. Since the levels of *gfp* expression in the AC that we achieved from the transgene appeared, based on assessment of GFP signal, lower than those seen in animals with endogenously *gfp*-tagged *lin-29a* (data not shown), we may speculate that some residual defect in the AC may prevent a fully penetrant phenotype. Moreover, additional tissues of expression may be relevant for either *lin-29a* or *mab-10*. This issue notwithstanding, our findings reveal an important role of MAB-10 and LIN-29a as effectors of *let-7* – LIN41 in its function to ensure vulval integrity, and they explain why *lin-29* and *let-7* mutations were previously found to yield incompatible phenotypes (Ecsedi et al., 2015).

What are then the events that fail and thereby cause vulval bursting when *lin-29a* and *mab-10* are not properly upregulated in *let-7* mutant animals? No obvious defects in vulval or uterine development of *let-7(PM)* animals could be detected (Ecsedi et al., 2015). This could suggest that a *let-7* function in a tissue other than the vulva or uterus is crucial for vulval integrity. However, previous work also revealed that in *let-7(PM)* animals, re-expression of *let-7* in epidermis, uterus and vulva sufficed to prevent vulval bursting, while re-expression in the epidermal hyp7 syncytium only was sufficient to restore some degree of epidermal differentiation but was insufficient to prevent vulval bursting (Ecsedi et al., 2015). Hence, although lack of suitably specific expression tools prevented a more refined dissection of the spatial requirements of *let-7* expression, it is likely that expression of *let-7* in the vulva, uterus and/or epidermal seam is required for vulval integrity. Therefore, it seems possible that attachments of uterine and/or vulval cells to each other and/or to the lateral seam are defective in *let-7* mutant animals. We expect that a detailed analysis of the expression patterns of *lin-29a* and *mab-10*, and the specific tissues and cell-types that require *let-7* expression for vulval integrity may help to define where *let-7 –* LIN41 regulate *lin-29a* and *mab-10* expression to prevent vulval rupturing. Future studies aimed at uncovering the transcriptionally regulated target genes of LIN-29a+MAB-10 may additionally provide insight into the underlying defects, and this and the study of MAB-3+DMD-3 targets will illuminate how *let-7 –* LIN41 direct cell fate and morphological changes at the transition to adulthood.

Our study focused on the functions of the LIN28 – *let-7* – LIN41 pathway in controlling self-renewal and transition to adulthood in *C. elegans*, yet these functions may be phylogenetically conserved. Specifically, the mammalian homologs of LIN28, *let-7* and LIN41 are known to be involved in the control of self-renewal versus differentiation programs in different cell types such as embryonic stem cells or neural progenitor cells (Faunes and Larrain, 2016). EGR and NAB proteins, homologous to the LIN41 targets MAB-10 and LIN-29a that we have shown here to be essential for these programs in *C. elegans* seam cells, have not yet been mechanistically linked to this pathway. However, they are crucial regulators of proliferation and/or terminal differentiation programs in different mammalian cell types, including embryonic stem cells, different blood cells or Schwann cells (Du et al., 2014; Laslo et al., 2006; Le et al., 2005; Min et al., 2008; Nguyen et al., 1993; Topilko et al., 1994; Worringer et al., 2014). Moreover, EGR1 overexpression antagonizes somatic cell reprogramming promoted through LIN41, although we note that EGR1 does not appear to be the closest homologue of LIN-29a (Pereira et al., 2018).

Yet more intriguingly, timing defects in human puberty have been linked to genetic variations in LIN28 (Faunes and Larrain, 2016), and in mice, both *Lin28(gf)* and *let-7(lf)* mutations were reported to delay the onset of puberty (Corre et al., 2016; Zhu et al., 2010). Whether mammalian LIN41 is involved in regulating the timing of puberty remains to be tested, and gene expression changes that control puberty remain poorly understood. However, we note that homologs of the transcriptional regulators downstream of LIN41 have known roles in the development of sex-specific structures in other animals including mammals. For example, mice homozygous for a null mutant of EGR1 are sterile, with females having an abnormal development of the ovary (Topilko et al., 1998). Moreover, DM domain-containing transcription factors control sexual differentiation across evolution and have crucial roles in the development of mammalian testis and germline (Kopp, 2012; Zhang and Zarkower, 2017). Given these tantalizing hints and apparent similarities, we suggest that it will be relevant to study whether LIN28 and *let-7* control the timing of mammalian sexual organ development through LIN41 and EGR/NAB or DM domain-containing transcription factors.

## ACKNOWLEDGEMENTS, funding

We thank Iskra Katic and Jun Liu for generating the *lin-41(bch28 xe70)* balancer allele, Sarah Carl for help with computational analysis of RIP-Seq data, Monika Fasler and Lan Xu for help with generating strains and Benjamin Towbin, Oliver Hobert, Laura Pereira, Iskra Katic, Chiara Azzi and Jun Liu for critical feedback on the manuscript. We thank Adam Mason and Douglas Portman for providing strains. Some strains were provided by the CGC, which is funded by NIH Office of Research Infrastructure Programs (P40 OD010440). This project has received funding from the Swiss National Science Foundation (#31003A_163447), the European Research Council (ERC) under the European Union’s Horizon 2020 research and innovation program (grant agreement n° 741269), and FMI core funding through the Novartis Research Foundation (to H.G.).

## AUTHOR CONTRIBUTIONS

F.A. and H.G. conceived the project. F.A., M.R. and H.G. designed experiments. F.A. executed most experiments. A.N. examined male tail and utse formation defects and generated strains. M.R. performed LIN41 RIP-Seq and generated strains. All authors analyzed data. F.A. and H.G wrote the manuscript.

## METHODS

### Generation of novel *lin-29*, *lin-29a*, *mab-10* and *mab-3* null mutant alleles using CRISPR-Cas9

Wild-type worms were injected with a mix containing 50 ng/μl pIK155, 100 ng/μl of each pIK198 with a cloned sgRNA template, 5 ng/μl pCFJ90 and 5 ng/μl pCFJ104, as previously described (Katic et al., 2015). Single F1 progeny of injected wild-type worms were picked to individual plates and the F2 progeny screened for deletions using PCR assays. After analysis by DNA sequencing, the alleles were outcrossed three times to the wild-type strain.

In order to obtain mutant alleles for *lin-29*, *mab-10* and *mab-3* that are undoubtedly null for the encoded protein, we injected two sgRNAs per gene to generate large deletions spanning almost the entire coding region. The following pairs were used: i) *lin-29* sgRNA1: gctggaaccaccactggctc, *lin-29* sgRNA2: atattatttatcagtgattg. ii) *mab-10* sgRNA1: gatgatgatgatgaagaggt, *mab-10* sgRNA2: gctcccggaatcttgaagct. iii) *mab-3* sgRNA1: aggagctctaatgctcaccg, *mab-3* sgRNA2: agctcagctcaatttgggcg.

For *lin-29*, we generated a large deletion spanning exons 2-11 and thus most of the coding region of both *lin-29a* and *lin-29b*. Specifically, the resulting *lin-29(xe37)* allele is a 14,801 bp deletion with a 2 bp insertion with the following flanking sequences: 5’ ggactctggaatagctggaa – *xe37* deletion – *xe37* insertion (aa) – aatatgaaaaatcattccta 3’. Translation of *xe37* yields only a short stretch of 28 amino acids (MDQTVLDSAFNSPVDSGIAG-KNMKNHSY*), containing the N-terminal 20 and the C-terminal 7 amino acids of LIN-29a. The small insertion leads to translation of an additional lysine (K).

For *mab-10*, the resulting *mab-10(xe44)* allele spans exons 3-9 and is a 2901 bp deletion with a 4 bp insertion with the following flanking sequences: 5’ ttatcatctcttacaactca – *xe44* deletion – *xe44* insertion (ctct) – tattttttgttttcctcg<u>tga</u> 3’. Translation of *xe44* yields a 58 amino acid stretch (MSSSSSSSLPTSSASTTTSSITSRPSASHHLESILSSSSSSPSILSSLTT-HSYFLFSS*) containing the N-terminal 50 amino acids of MAB-10, followed by 8 additional amino acids, translated from the small insertion and the *mab-10* 3’UTR, and a stop codon (underlined in the flanking sequence above).

For *mab-3*, the resulting *mab-3(xe49)* allele is a 4291 bp deletion starting in exon 2, just downstream of the ATG start codon of the longer isoform, and ending downstream of the stop codon. The flanking sequences are: 5’ tttgcagaggagctctaatg – *xe49* deletion – ctccgcccacactttcccag 3’. Translation of *xe49* is presumably initiated at the normal ATG start codon, but then translates a sequence that is normally non-coding, yielding a stretch of 31 amino acids (MLRPHFPRITVLFLALRLSFSFPLSLFYLGK*) unrelated to the MAB-3 protein.

In order to specifically mutate *lin-29a* without affecting expression of *lin-29b*, we deleted part of the coding exons specific to *lin-29a*, at the same time introducing a frame-shift in the downstream *lin-29a* reading frame. By injecting two sgRNAs (sgRNA1: gctggaaccaccactggctc, sgRNA2: gtggcaggagagaattctga), we obtained the *lin-29a(xe40)* allele, a 1102 bp deletion covering exons 2-4, introducing a frame-shift in the *lin-29a* reading frame with a predicted stop codon in exon 6. The deletion has the following flanking sequences: 5’ ctctggaatagctggaaccac – *xe40* deletion – attctctcctgccacatcat 3’. Translation of *xe40* yields a protein with the N-terminal 22 amino acids of LIN-29A (MDQTVLDSAFNSPVDSGIAGTT), followed by a stretch of 69 out-of-frame amino acids and a stop codon.

### Isoform-specific GFP::3xFLAG tagging of endogenous *lin-29a* using CRISPR-Cas9

In order to specifically tag *lin-29a* at the N-terminus, the following mix was injected into wild-type worms (Dickinson et al., 2015; Katic et al., 2015): 50 ng/μl pIK155, 100 ng/μl of pIK198 with a cloned sgRNA template (atattatttatcagtgattg), 2.5 ng/μl pCFJ90, 5 ng/μl pCFJ104 and 10 ng/μl pDD282 with cloned homology arms. Recombinants were isolated according to the protocol by Dickinson *et al*. (Dickinson et al., 2015), verified by DNA sequencing and outcrossed three times. The plasmid for homologous recombination, pFA27, was prepared by restriction digest of pDD282 with ClaI and SpeI, followed by a Gibson assembly reaction (Gibson et al., 2009) with two gBlocks^®^ Gene Fragments (Integrated DNA Technologies) (Table S5).

### Generation of a balancer allele for *lin-41(xe8)* using CRISPR-Cas9

The *lin-41(xe8)* allele is not temperature-sensitive like *let-7(n2853)* and therefore causes lethality at any temperature (Ecsedi et al., 2015). In order to maintain *lin-41(xe8)* animals, a balancer null allele, *lin-41(bch28)*, was previously created by inserting an expression cassette driving ubiquitous nuclear GFP from the *eft-3* promoter into the *lin-41* coding sequence (Katic et al., 2015). To avoid generating a wild-type *lin-41* copy – and a recombined *lin-41(bch28 xe8)* allele – through intragenic recombination of *lin-41(bch28)* with *lin-41(xe8)*, we additionally deleted a large part of the *lin-41* coding sequence together with the part of the *lin-41* 3’UTR containing the *let-7* complementary sites within the *lin-41(bch28)* allele. To this end, *lin-41(bch28)* heterozygous worms were injected with a mix containing 50 ng/μl pIK155, 100 ng/μl of each pIK198 with a cloned sgRNA, 5 ng/μl pCFJ90 and 5 ng/μl pCFJ104, as previously described (Katic et al., 2015). We injected two plasmids encoding sgRNAs, sgRNA1 (ggtgactgaatcattgacgg) and sgRNA2 (agaaggtttcaatggttcag), cutting in the third coding exon and the 3’UTR of *lin-41*, respectively. Single F1 progeny of injected wild-type worms were picked to individual plates and the F2 progeny were screened for expected deletions in *lin-41(bch28)* by PCR. The *lin-41(bch28 xe70)* allele thus obtained was further validated by DNA sequencing and outcrossed three times to the wild-type strain before crossing it with *lin-41(xe8)* heterozygous animals. The final *lin-41(bch28 xe70)* allele consists of the inserted expression cassette, as described in (Katic et al., 2015), followed by an additional deletion of the region with the following flanking sequences: 5’ ggctcactatttgacactcc – *xe70* deletion (6395 bp) – accattgaaaccttctccc 3’.

### C. elegans

Worm strains used in this study are listed in Table S6. Bristol N2 was used as the wild-type strain. To obtain arrested L1 larval stage worms, embryos were extracted from gravid adults using a bleaching solution (30% (v/v) sodium hypochlorite (5% chlorine) reagent (Thermo Fisher Scientific; 419550010), 750 mM KOH). Next, the embryos and subsequently hatched L1 larvae were incubated overnight in the absence of food, at room temperature in M9 buffer (42 mM Na_2_HPO_4_, 22 mM KH_2_PO_4_, 86 mM NaCl, 1 mM MgSO_4_). If not otherwise specified, worms were grown on 2% NGM agar plates with *Escherichia coli* OP50 bacteria (Stiernagle, 2006). For RIP-Seq experiments, worms were grown on enriched peptone plates with *Escherichia coli* NA22 bacteria (Evans, 2006). For RNAi experiments, arrested L1s were plated on RNAi-inducing NGM agar plates with *Escherichia coli* HT115 bacteria containing plasmids targeting the gene of interest (Ahringer, 2006).

### Construction of transgenic single-copy GFP reporters

Transgenes were cloned into the the destination vector pCFJ150 (Frokjaer-Jensen et al., 2008) using the MultiSite Gateway Technology (Thermo Fisher Scientific) as described in Table S5. Worm lines with integrated transgenes were obtained by single-copy integration into chromosome II (ttTi5605 locus), using a protocol for injection with low DNA concentration (Frøkjær-Jensen et al., 2012).

### Confocal imaging

For confocal imaging of the *mab-3* and *dmd-3* reporter worm lines (HW1803, HW1798, HW1827 and HW1828), synchronized arrested L1 stage larvae were grown for 20 hours at 25 °C on RNAi-inducing plates with HT115 bacteria. The bacteria either contained the L4440 parental RNAi vector without insert (denoted “mock RNAi”) or with an insert targeting *lin-41* (Fraser et al., 2000). For confocal imaging of endogenously tagged GFP::LIN-29a (HW1826 and HW1882), worms were grown at 25 °C on OP50 bacteria. Worms were imaged on a Zeiss LSM 700 confocal microscope driven by Zen 2012 Software after mounting them on a 2% (w/v) agarose pad with a drop of 10 mM levamisole solution. Differential Interference Contrast (DIC) and fluorescent images were acquired with a 40x/1.3 oil immersion objective (1024×1024 pixels, pixel size 156nm). Using the Fiji software (Schindelin et al., 2012), images were processed after selecting representative regions. Worms of the same worm line were imaged and processed with identical settings.

### Seam cell imaging and quantification

Arrested L1 larvae were plated on OP50 bacteria and synchronized worms were grown at 25 °C for 36-38 hours (late L4 stage) or 40-42 hours (young adult stage), with the exact developmental time assessed by staging of individual worms according to gonad length and vulva morphology. Worms were mounted to a 2 % (w/v) agarose pad and immobilized in 10 mM levamisole. Fluorescent and Differential Interference Contrast (DIC) images were acquired with a Zeiss Axio Observer Z1 microscope using the AxioVision SE64 software. Selections of regions and processing of images was performed with Fiji (Schindelin et al., 2012). Seam cell quantifications were performed by counting all clearly visible fluorescent cells expressing an *scm::gfp* transgene (Koh and Rothman, 2001) of the upper lateral side in mounted worms. For worm lines containing mnC1-balanced animals, *myo-2p*::GFP-positive animals were excluded from imaging and quantifications. To score *lin-41(xe8)* homozygous animals within a population of balanced *lin-41(xe8)*/*lin-41(bch28 xe70)* animals, all *eft-3p*::*gfp::h2b* expressing (i.e., balancer carrying) animals were excluded from the analysis.

### Imaging and quantification of male tail phenotypes

After obtaining arrested L1 larvae, synchronized populations of *him-5(e1490)* mutant worms in the different genetic backgrounds were grown on OP50 bacteria at 25 °C. After growing males to early, mid or late L4 larvae as well as to young adults, they were mounted on a 2 % (w/v) agarose pad and immobilized in 10 mM levamisole for imaging and quantification of the tail phenotypes with a Zeiss Axio Observer Z1 microscope. Differential Interference Contrast (DIC) images were acquired using the AxioVision SE64 software. Selections of tail regions and processing of images was performed with Fiji (Schindelin et al., 2012).

To quantify tail retraction defects, male tail phenotypes were scored in mounted worms at the late L4 and at the young adult stage. At the late L4 stage, tail retraction defects were scored as unretracted (no cell retraction visible), partially retracted (cells less retracted compared to wild-type worms at the same developmental stage) or over-retracted tails (further cell retraction compared to wild-type worms at the same developmental stage). At the young adult stage, abnormal tails were categorized into over-retracted, Lep and unretracted (very long spike, compromised rays and fan structures) phenotypes. Tails with a spike extending beyond the fan were counted as “Lep” and tails with smaller spikes were considered wild-type (as in our hands, those were difficult to distinguish from wild-type tails). While most *lin-41(bx37)* or *lin-41(bx42)* mutant males had a Lep tail, all *lin-41(xe8)* animals showed the distinct and more severe “unretracted” phenotype. 85 % of *let-7(n2853)* mutant males also displayed a completely unretracted tail. In 15 % of the scored *let-7(n2853)* mutant males, the tail cell started retracting at the time point of analysis (young adult stage), still resulting in a similarly elongated tail, but with a rounded tip. This phenotype was categorized as “unretracted”, as it was more similar to fully unretracted tails than to Lep tails. The observed delayed start of the retraction program may be due to remaining *let-7* activity in the *let-7(n2853)ts* mutant. For worm lines containing mnC1-balanced or *lin-41(bch28 xe70)*-balanced animals, GFP-positive animals were excluded from the analysis.

### Quantification of vulval bursting phenotypes

Arrested L1 larvae were plated on NGM agar plates with OP50 bacteria and grown as synchronized populations at 25 °C for 45 hours. This developmental stage corresponded to an adult stage approximately five hours after the first *let-7(n2853)* or *lin41(xe8)* animals burst through their vulva. Using a dissecting microscope, worm phenotypes were scored directly on the NGM agar plates by distinguishing animals with part of the intestine protruding through the vulva (“burst”) from other animals (“non-burst”). For each genotype, at least 400 worms were scored in each experiment. For worm lines containing mnC1-balanced or *lin-41(bch28 xe70)*-balanced animals, GFP-positive animals (distinguishable from unbalanced worms using a GFP-filter on the dissecting microscope) were excluded from the analysis.

### Imaging and quantification of uterine-vulval connection phenotypes

Synchronized populations of arrested L1 larvae were plated on OP50 bacteria and grown to late L4 stage larvae at 25 °C for 36-38 hours. Worms were mounted to a 2 % (w/v) agarose pad, immobilized in 10 mM levamisole and their vulval-uterine regions imaged with a Zeiss Axio Observer Z1 microscope using Differential Interference Contrast (DIC) and the AxioVision SE64 software. Vulval-uterine regions for representative images were selected and processed with the Fiji software (Schindelin et al., 2012). Phenotypes of mounted worms were counted and assigned to two categories, worms with a normal thin utse structure and worms with an abnormal thicker cell layer. For worm lines containing mnC1-balanced individuals, *myo-2p*::GFP-positive animals were excluded from the quantifications.

### RNA co-immunoprecipitation coupled to RNA-sequencing (RIP-seq)

RNA co-immunoprecipitations (RIPs) were performed with semi-synchronous L3/L4 stage populations of *lin-41(n2914); him-5(e1490)* mutant worms expressing *flag::gfp::lin-41* (Aeschimann et al., 2017) and of *him-5(e1490)* mutant worms expressing *flag::gfp::sart-3* (Rüegger et al., 2015) as a control, following the protocol of our previous publication (Aeschimann et al., 2017). For each RIP, the RNA bound to the beads as well as a sample of input RNA were extracted using Tri Reagent (Molecular Research Center; TR 118) according to the manufacturer’s recommendations. The input RNA samples were diluted to match the low RNA concentrations of 5-10 ng/μl of the immunoprecipitated RNA. Libraries were prepared with the TruSeq Stranded mRNA HT Sample Prep Kit (Illumina; RS-122-2103) and 50-bp single end reads were sequenced on an Illumina HiSeq2000 machine.

### RIP-seq analysis

PCR duplicates were first removed by collapsing reads with an identical 5’ end coordinate to a single read. De-duplicated reads were then aligned to the May 2008 (ce6) C. elegans genome assembly from UCSC (Rosenbloom et al., 2015). Alignments were performed using the qAlign function from the QuasR R package (v. 1.20.0) (Gaidatzis et al., 2015), with the reference genome package (“BSgenome.Celegans.UCSC.ce6”) downloaded from Bioconductor (https://www.bioconductor.org) and with the parameter “splicedAlignment=TRUE”, which calls the SpliceMap aligner with default parameters (Au et al., 2010). The resulting alignments were converted to BAM format, sorted and indexed using Samtools (version 1.2) (Li et al., 2009). Gene coverage was quantified using annotations downloaded from WormBase (version WS190; ftp://ftp.wormbase.org/pub/wormbase/releases/WS190/). Reads overlapping all annotated exons for each gene were counted. For plotting, samples were normalized by the mean number of counts mapping to exons in all samples and then log_2_-transformed after adding a pseudocount of 8. To determine genes significantly bound by LIN-41, a model was constructed using edgeR (v. 3.22.3) (Robinson et al., 2010) containing a term for sequencing batch, library type (IP or input) and protein (LIN-41 or SART-3), as well as an interaction term between library type and protein. Testing for significance of the interaction term with a likelihood ratio test identified genes for which the enrichment of IP vs. input was significantly greater for LIN-41 than for the SART-3 control. After conducting a multiple hypothesis test correction, we applied a cutoff of FDR < 0.05 to determine a final set of bound genes. All computations were performed using R (v. 3.5.1).

### Data availability

All RIP-seq data generated in this study have been deposited in the NCBI Gene Expression Omnibus (Edgar et al., 2002) under GEO Series accession number GSE120405.

## SUPPLEMENTAL INFORMATION

**Figure S1.**
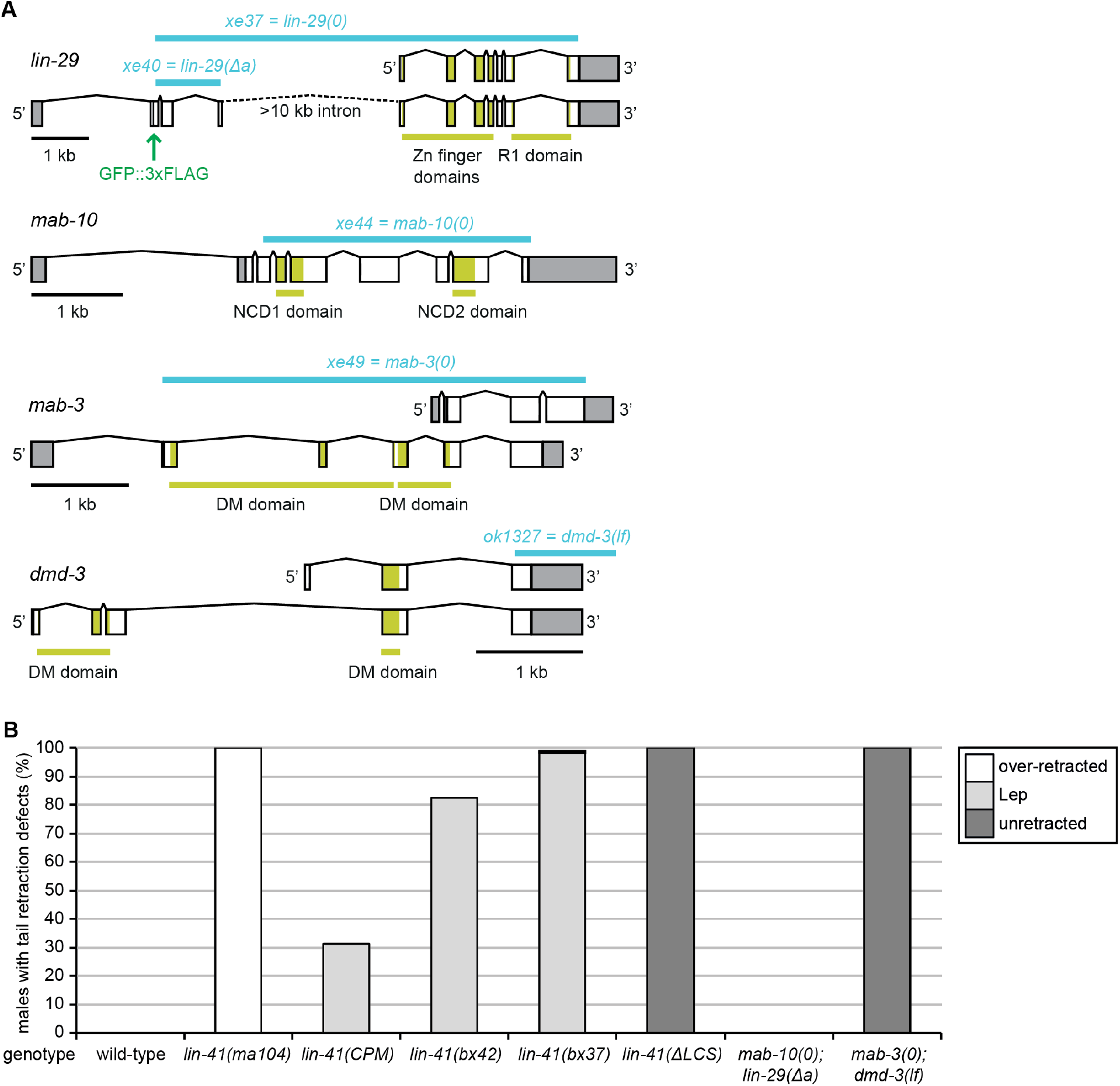
Novel mutant alleles and male tail phenotype experiments. (A) Illustration of the mutant alleles (in blue) used in this study to disrupt the function of LIN41 target genes. The position of the endogenous GFP::3xFLAG inserted at the N-terminus of LIN-29a (*lin-29(xe63)*) is indicated in green. (B) Quantification of the male tail phenotypes at the young adult stage of the indicated genotypes. Shown are the percentages of animals with over-retracted, Lep or unretracted tails. n≥100, except for *lin-41(ma104)* (n=55), which displays an over-retracted male tail phenotype and was included as a control. Worms were grown at 25 °C.

**Table S1.**
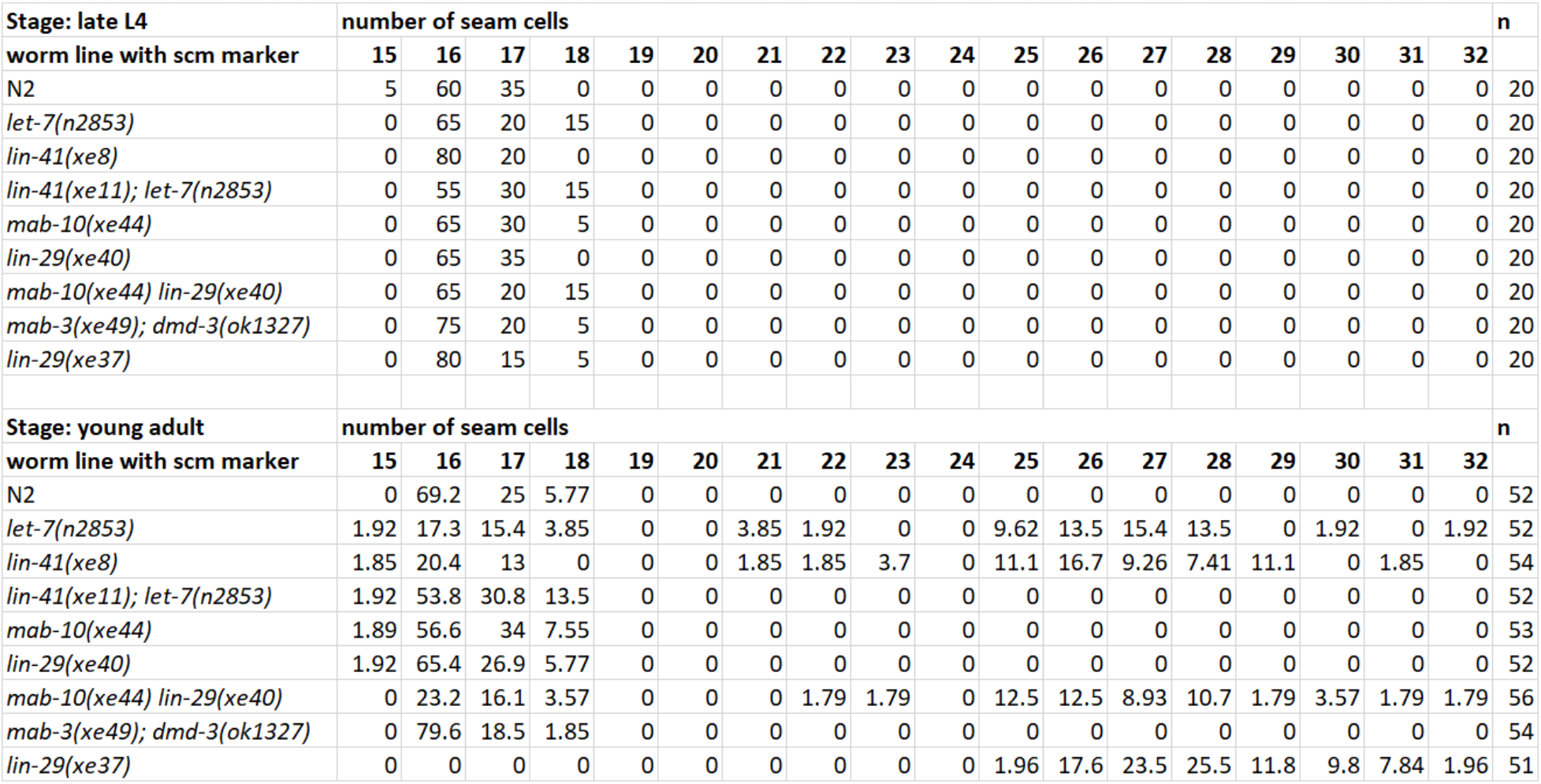
Number of seam cells in late L4 and young adult stages. (numbers are given in percentage of animals, n = number of scored animals)

**Table S2.** Male tail retraction phenotypes. (numbers are given in percentage of animals, n = number of scored animals)

|  |  |  |  |  |  |
| --- | --- | --- | --- | --- | --- |
| **Stage: late L4** |  |  |  |  |  |
| **worm line (in him-5(e1490) background)** | **Wild-type\*\*\*** | **Partial retraction** | **Unretracted** | **Over-retracted** | **n** |
| N2 (wild-type) | 100 | 0 | 0 | 0 | 109 |
| *let-7(n2853)* | 0 | 3.76 | 96.24 | 0 | 133 |
| *lin-41(xe8)\** | 0 | 0 | 100 | 0 | 103 |
| *lin-41(xe11)* | 58.12 | 38.46 | 3.42 | 0 | 117 |
| *lin-41(xe11); let-7(n2853)* | 98.59 | 0.70 | 0.70 | 0 | 142 |
| *lin-29(xe40) mab-10(xe44)\*\** | 98.17 | 1.83 | 0 | 0 | 109 |
| *mab-3(xe49); dmd-3(ok1327)* | 0 | 0 | 100 | 0 | 108 |
| *lin-41(bx37)* | 0 | 87.5 | 12.5 | 0 | 104 |
| *lin-41(bx42)* | 0 | 96.75 | 3.25 | 0 | 123 |
| *lin-41(ma104)* | 7.5 | 0 | 1.25 | 91.25 | 80 |
| **Stage: young adult** |  |  |  |  |  |
| **worm line (in him-5(e1490) background)** | **Wild-type\*\*\*** | **Lep (outside)** | **Unretracted** | **Over-retracted** | **n** |
| N2 (wild-type) | 100 | 0 | 0 | 0 | 100 |
| *let-7(n2853)* | 0 | 0 | 100**** | 0 | 133 |
| *lin-41(xe8)\** | 0 | 0 | 100 | 0 | 105 |
| *lin-41(xe11)* | 68.78 | 31.22 | 0 | 0 | 189 |
| *lin-41(xe11); let-7(n2853)* | 100 | 0 | 0 | 0 | 100 |
| *lin-29(xe40) mab-10(xe44)\*\** | 100 | 0 | 0 | 0 | 117 |
| *mab-3(xe49); dmd-3(ok1327)* | 0 | 0 | 100 | 0 | 106 |
| *lin-41(bx37)* | 0.88 | 98.25 | 0.88 | 0 | 114 |
| *lin-41(bx42)* | 17.43 | 82.57 | 0 | 0 | 109 |
| *lin-41(ma104)* | 0 | 0 | 0 | 100 | 55 |
| *these worms were obtained from a balanced line: *lin-41(xe8)/lin-41(bch28 xe70)* |  |  |  |  |  |
| **these worms were obtained from a balanced line: *lin-29(xe40) mab-10(xe44) / mnC1* |  |  |  |  |  |
| ***weak Lep phenotypes (spike inside the fan) were considered wild-type |  |  |  |  |  |
| ****in 20 out of the 133 worms, the tail cells started retracting at this time point of analysis. |  |  |  |  |  |

**Table S3.** Vulval bursting phenotypes. (numbers are given in percentage of burst animals in each of three independent experiments (exp) with n ≥ 400 scored animals)

| **Table 3A** |  |  |  |  |  |
| --- | --- | --- | --- | --- | --- |
| **worm line** | **% burst exp1** | **% burst exp2** | **% burst exp3** | **AVERAGE** | **STDEV** |
| wild-type (N2) | 0 | 0 | 0 | 0 | 0 |
| *let-7(n2853)* | 92.02 | 90.71 | 85.27 | 89.33 | 3.58 |
| *lin-41(xe8)* | 86.86 | 89.93 | 88.76 | 88.51 | 1.55 |
| *lin-41(xe11); let-7(n2853)* | 0 | 0 | 0 | 0 | 0 |
| *lin-29(xe40)* | 1.69 | 1.58 | 2.12 | 1.80 | 0.28 |
| *mab-10(xe44)* | 0.23 | 0 | 0.24 | 0.16 | 0.13 |
| *mab-10(xe44) lin-29(xe40)* | 15.35 | 17.01 | 9.98 | 14.11 | 3.68 |
| *mab-3(xe49); dmd-3(ok1327)* | 0.22 | 0 | 0 | 0.07 | 0.13 |
| *mab-3(xe49) mab-10(xe44) lin-29(xe40); dmd-3(ok1327)* | 14.02 | 15.56 | 10.30 | 13.29 | 2.71 |
| *lin-29(xe37)* | 2.34 | 0.96 | 2.07 | 1.79 | 0.73 |
| *mab-10(xe44) lin-29(xe37)* | 1.91 | 1.20 | 1.56 | 1.56 | 0.36 |
| **Table 3B** |  |  |  |  |  |
| **worm line** | **% burst exp1** | **% burst exp2** | **% burst exp3** | **AVERAGE** | **STDEV** |
| *let-7(n2853)* | 90.65 | 96.12 | 94.31 | 93.69 | 2.79 |
| *lin-29(xe37); let-7(n2853)* | 0.71 | 0.99 | 0.22 | 0.64 | 0.39 |
| *lin-29(xe40); let-7(n2853)* | 20.21 | 12.47 | 13.08 | 15.25 | 4.30 |
| *mab-10(xe44); let-7(n2853)* | 93.67 | 91.51 | 97.44 | 94.20 | 3.00 |
| *mab-10(xe44) lin-29(xe40); let-7(n2853)* | 15.99 | 15.02 | 13.33 | 14.78 | 1.35 |
| **Table 3C** |  |  |  |  |  |
| **worms from lines balanced with mnC1** | **% burst exp1** | **% burst exp2** | **% burst exp3** | **AVERAGE** | **STDEV** |
| *mab-3(xe49) mab-10(xe44) lin-29(xe40); dmd-3(ok1327)* | 8.29 | 12.69 | 16.14 | 12.38 | 3.94 |
| *mab-10(xe44) lin-29(xe40)* | 13.77 | 14.55 | 20.99 | 16.44 | 3.96 |
| *xeSi417 mab-10(xe44) lin-29(xe40)* | 51.75 | 50.64 | 50.84 | 51.08 | 0.59 |
| *lin-29(xe37)* | 0.68 | 0.98 | 0.92 | 0.86 | 0.16 |
| *xeSi417 lin-29(xe37)* | 95.12 | 92.84 | 93.32 | 93.76 | 1.20 |

**Table S4.**
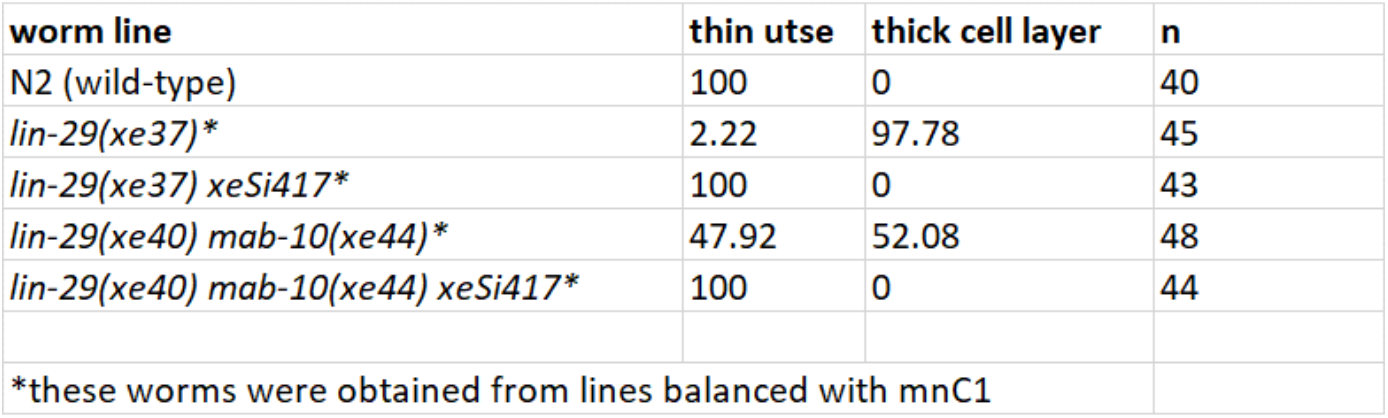
Presence of the thin utse structure or thick cell layers between vulva and uterus. (numbers are given in percentage of animals, n = number of scored animals)

**Table S5.**
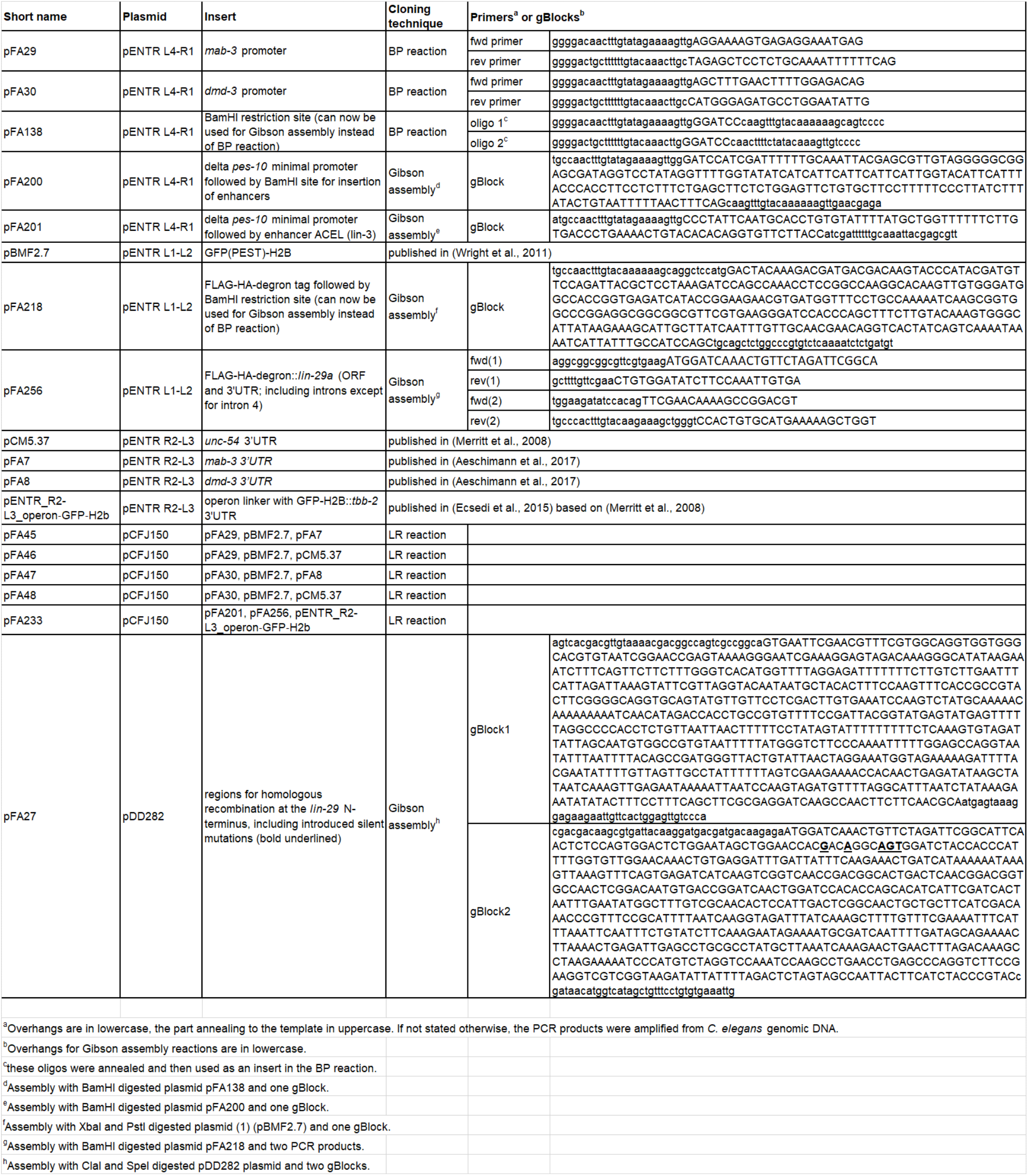
Plasmids used in this study.

**Table S6:**
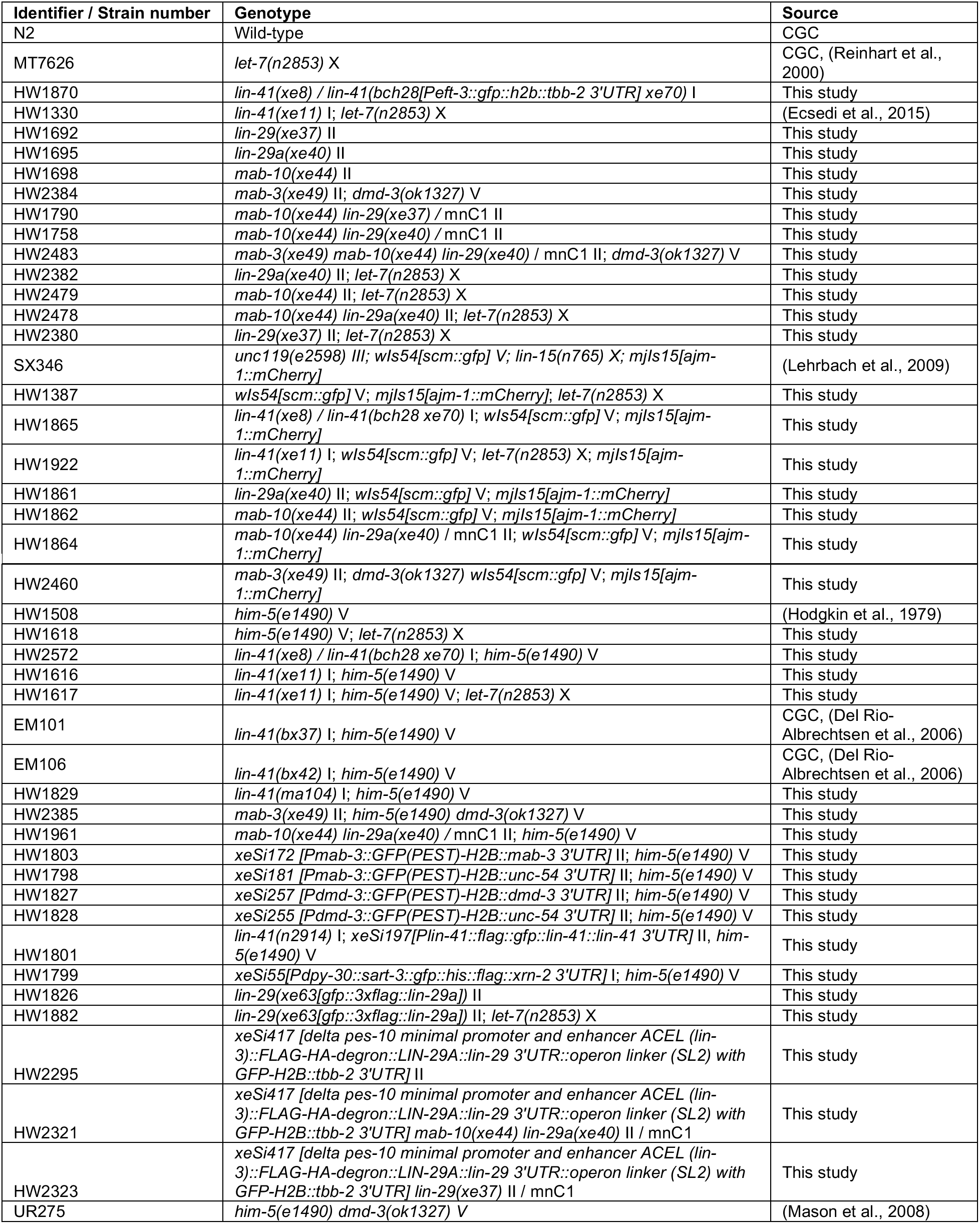
Worm strains used in this study.

